# Strong but diffuse genetic divergence underlies differentiation in an incipient species of marine stickleback

**DOI:** 10.1101/2025.09.19.677379

**Authors:** Alexandra Sumarli, Colby Behrens, Gina Lucas, Michael J. Olufemi, Paul Bentzen, Alison M. Bell, Frederic J.J. Chain, Kieran Samuk

**Affiliations:** Department of Evolution, Ecology, and Organismal Biology, University of California, Riverside, Riverside, California, USA; Department of Evolution, Ecology and Behavior, School of Integrative Biology, University of Illinois at Urbana-Champaign, Urbana, IL, USA; Program in Ecology, Evolution and Conservation, University of Illinois Urbana-Champaign, Urbana, IL, USA; Carl R. Woese Institute for Genomic Biology, University of Illinois Urbana-Champaign, Urbana, IL, USA; Program in Neuroscience, University of Illinois Urbana-Champaign, Urbana, IL, USA; Department of Biological Sciences, University of Massachusetts Lowell, Lowell, Massachusetts, USA; Department of Biology, Dalhousie University, Halifax, NS, Canada

**Keywords:** Reproductive Isolation, gene flow, stickleback, Copy Number Variation, incipient species

## Abstract

Understanding how lineages proceed along the “speciation continuum” and how species boundaries are maintained over time remain central questions in evolutionary biology. Populations early in the speciation process can give us detailed insight into the reproductive barriers that first initiate speciation. In this study, we explore the nature of genomic divergence between two sympatric marine stickleback ecotypes from Atlantic Canada, “whites” and “commons”. Males of each ecotype exhibit distinct nuptial colorations, nesting habits, and parental care strategies. Using population genomic analyses of SNPs and copy number variants (CNVs; deletions and duplications) we show that whites and commons consistently form distinct populations. We uncover genomic differentiation in the white ecotype characteristic of an incipient species, showing extremely low genome-wide differentiation (F_ST_) and very recent divergence (∼1 kya). Demographic analysis detected very low levels of ongoing gene flow between populations. Our results and prior genomic studies suggest that reproductive isolation is being maintained between ecotypes despite recent evidence that hybridization in nature does occur. Contrary to other systems, we found many small, but dispersed regions of high differentiation throughout the genome rather than explicitly within chromosomal inversions or the sex chromosomes. On chromosomes VII and XVI, we identified CNVs overlapping genes enriched for olfaction, which may play a role in differences in reproductive strategies between ecotypes. Ultimately, our results demonstrate that genome-wide rather than localized differences can underlie the early stages of divergence, and that this pattern is corroborated by both SNPs and CNVs.

## INTRODUCTION

Speciation, the evolution of reproductive isolation (RI) between populations, is one of the most fundamental and historically contentious areas in evolutionary biology. A central goal of contemporary speciation research is to understand the evolutionary mechanisms and specific isolating barriers involved in reducing gene flow between lineages (e.g., Ravinet et al., 2017; Schluter and Rieseberg, 2022; Stankowski et al., 2024). The pervasiveness of gene flow between divergent lineages in nature raises many questions about the genetic basis of RI. Namely, are there specific “speciation genes” (i.e., genes that directly contribute to RI; Blackman, 2016) or genomic regions underlying RI, what are the relative impacts of pre- and postzygotic barriers on restricting gene flow, and what evolutionary forces are responsible for promoting or inhibiting diversification? While there is a wide body of existing theoretical and empirical work in the field of speciation (e.g., Coyne and Orr, 2004; Coughlan and Matute, 2020; Thompson et al., 2024), we still lack a unified theoretical framework for understanding the genomic basis of RI.

Populations in the early stages of divergence (i.e., incipient species) can offer key insights into how RI originates and accumulates over time (e.g., sunflower ecotypes, Andrew and Riesberg, 2013; threespine stickleback from Jordeweiher pond, Marques et al., 2016; southern capuchino seedeaters, Turbek et al., 2021). Traditionally, speciation gene studies focused on “good” species or populations that have been isolated for longer evolutionary timescales (reviewed in Presgraves 2010). A limitation of this approach is that it is retrospective: the genes responsible for initiating divergence have already been obscured by the genetic differences that accrued after RI was completed (Via 2009). Instead, incipient species that exhibit partial RI are uniquely suited for identifying the genes that are directly involved in causing rather than maintaining population divergence.

Many hypotheses have proposed that speciation genes should be localized in regions of suppressed recombination, such as within chromosomal inversions (Kirkpatrick and Barton, 2006; Rieseberg, 2001). Inversions play a role in RI by inhibiting gene flow and preserving favorable combinations of alleles that contribute to speciation and local adaptation. While inversions have garnered considerable attention in recent years (e.g., *Helianthus* sunflowers: Huang, et al., 2020; threespine stickleback: Jones et al., 2012; *Timema* stick insects: Nosil et al., 2023), less is known about how other types of structural polymorphisms impact speciation. For example, copy number variations (CNVs) are deletions and duplications that result in different numbers of gene copies segregating among individuals. CNVs can alter phenotypes and consequently fitness through mechanisms including modifying the regulation, structure, and function of genes (reviewed in Iskow et al., 2012). As such, CNVs are known to promote population-specific adaptations such as toxin resistance in Atlantic killifish (Reid et al., 2016) and starch digestion in humans (Perry et al., 2007). They have also been linked to olfactory receptors potentially involved in assortative mating between divergent populations (pigs: Paudel et al., 2015; European house mice: North et al., 2020).

Like inversions, sex chromosomes are also invoked as hotspots for speciation genes (Dufresnes and Crochet, 2022; Qvarnström and Bailey, 2009). In part this is due to the distinct inheritance patterns of XY and ZW systems that result in the sex chromosomes harboring different effective population sizes (N_e_) and levels of genetic diversity (π) compared to autosomes (reviewed in Wilson Sayres, 2018). Under neutral conditions, the X chromosome (Z in ZW systems) has ¾ the N_e_ of the autosomes while the Y chromosome (W in ZW systems) has ¼ the N_e_ of the autosomes. Reduced N_e_ coupled with enhanced drift can cause sex chromosomes to evolve faster relative to autosomes (i.e., faster X or Z effects; Meisel and Connallon, 2013).

This can lead to relative genetic differentiation (F_ST_) being significantly higher on the sex chromosome compared to the autosomes as is seen in numerous bird species (e.g., Irwin, 2018; Schield et al., 2021). According to theory, sex chromosomes can further promote RI when genes underlying prezygotic isolation and mate preference (e.g., plumage coloration, courtship displays, sperm-egg proteins) accumulate and become linked on the sex chromosomes due to sexually antagonistic selection (Albert and Otto, 2005; Gavrilets, 2000).

Threespine stickleback (*Gasterosteus aculeatus,* hereafter “stickleback”) are a powerful system for studying behavior, adaptation, and speciation (reviewed in Reid et al 2021). Stickleback are ancestrally marine fish that have repeatedly colonized freshwater for at least the last 10-20 million years, most recently following the Last Glacial Maximum (LGM; Bell, 1994). These recent and repeated colonizations of different freshwater habitats have led to the existence of numerous coexisting ecotypes throughout their Holarctic range (McKinnon and Rundle, 2002). In benthic and limnetic species pairs, RI is mediated by strong divergent natural selection based on adaptation to different ecological niches (Schluter and McPhail, 1992) and assortative mating based on body size (Conte and Schluter, 2013). While much speciation research has been allocated towards stickleback ecotypes with distinct trophic morphologies or freshwater residents (e.g., benthic-limnetic, lake-stream), fully marine and anadromous sticklebacks (i.e., marine stickleback that breed in fresh or brackish water) have received far less attention. Furthermore, much focus has centered on relatively older populations from the Pacific and eastern Atlantic (e.g., the Pacific Northwest, Japan, Baltic Sea; Fang et al., 2018, 2020). The biology and evolutionary history of more recently derived sticklebacks from the western Atlantic remain underexplored (Haines, 2023).

The “white” threespine stickleback (*Gasterosteus aculeatus*) ecotype from Nova Scotia, Canada offers the exciting opportunity to investigate the genetic and phenotypic changes involved in the early stages of divergence. Male “whites” exhibit pearlescent-white nuptial coloration and often occur in sympatry with larger sized male “commons” during their breeding season when they spawn in brackish water lagoons (Blouw and Hagen, 1990). Compared to male commons, male whites exhibit distinct courtship displays, lack parental care behavior, and nest exclusively in filamentous algae (Behrens et al. 2024; Blouw, 1996; Jamieson et al., 1992ab). Female whites lay smaller eggs that can be easily dispersed compared to female commons which lay larger, tightly clumped eggs (Behrens et al., 2025a; Grant, 1993). Based on these phenotypic differences, it is thought that the white stickleback likely represents an incipient species (Blouw, 1996; Samuk, 2016).

The most comprehensive and recent genomic studies of this system, Samuk (2016) and Behrens et al. (in review, 2025b) found that whites and commons form discrete genetic clusters based on RADseq-derived SNPs. The former study examined 296 individuals and found that these two ecotypes exhibited extremely low genomic divergence (F_ST_) and that divergence with gene flow (i.e., under an IM model) between these lineages occurred recently (<12,000 ya). While white-common hybrid offspring in laboratory settings are viable and there are no intrinsic postzygotic barriers between ecotypes, prior studies suggest there is a scarcity of F1 hybrids in nature (Blouw, 1996; Samuk, 2016). Recently, Behrens et al (in review, 2025b) found two putative white-common hybrids based on random sampling of 200 sticklebacks from across the range of whites and commons. Behrens et al (in review, 2025b) also demonstrated that white-common hybrid fathers exhibited dysregulated parental care and proposed this as a possible mechanism for RI between ecotypes. Interestingly, whites do not exhibit significant ecological differences from commons based on stable isotopes (Samuk, 2016). Furthermore, evidence of assortative mating and partially non-overlapping breeding times is inconclusive (Blouw and Hagen, 1990; Corney and Weir, 2024). Taken together, the evolutionary forces and genomic patterns of divergence underlying this system remain unclear.

Here, we employ whole genome sequencing (WGS) in multiple populations of whites and commons to explore the nature of genomic divergence and elucidate their evolutionary history. Unlike Behrens et al. (in review, 2025b) and Samuk (2016), our study leverages CNV data alongside replicated whole genome data from multiple populations from each ecotype, and the most recent threespine stickleback reference genome, which includes an assembly of the Y chromosome (Nath et al., 2021; Peichel et al., 2020). We sought to answer three main questions:

First, do white and common ecotypes appear to be distinct populations across their range? Second, does the white stickleback indeed represent an incipient species? Lastly, are there specific regions of the genome that contribute to the discrete population identities of whites and commons? In answering these questions, we seek to gain a deeper understanding of how species boundaries are formed during the earlier stages of speciation.

## MATERIALS AND METHODS

### Sample collection and whole genome sequencing

In May of 2011, adult commons from Overton (43.83 N, 66.15 W), Nova Scotia, Canada were collected by Stanley King (Bentzen Lab, Dalhousie University, Halifax, NS) using minnow traps. After euthanasia, whole bodies were stored in 95% ethanol. In June and July of 2017 and 2018 during the field work conducted by Haley et al. (2019), fin clips from adult commons and whites from Rainbow Haven (44.65 N, 63.42 W), Canal Lake (44.50 N, 63.90 W), and Baddeck (46.10 N, 60.74 W), Nova Scotia were collected by Laura Weir, Anne Haley and Anne Dalziel (Saint Mary’s University, Halifax, NS). Because whites and commons can be found in sympatry at these three sites, morphology and breeding behaviors were observed for ∼ five to 10 minutes before capture. Male whites were identified by their bright white courtship coloration and smaller size relative to commons and caught via dip netting near their nests in filamentous algae. Male commons were caught by their nests in mud or sand. Gravid females were collected in minnow traps and assigned as white or common based upon size. After capture, a small caudal fin clip was taken from each fish and stored in 95% ethanol

In early June of 2019, 20 adult commons were collected from Cherry Burton Road, New Brunswick, Canada (46.00 N, 64.00 W) and 20 whites were collected from Canal Lake, Nova Scotia, Canada (44.5 N, 63.90 W) using unbaited mesh minnow traps or dip nets. 10 males and 10 females were collected at both sites. Fish were then transported to the University of Illinois in Urbana-Champaign, where they underwent behavioral testing before sacrificing for tissue collection.

For samples collected prior to 2019, we extracted DNA from 48 specimens following the protocol of Elphinstone et al., (2003). DNA samples were initially quantified using a Quant-iT PicoGreen dsDNA Assay kit (Invitrogen, Richardson USA) in 200 µL volumes using 1 µL of DNA sample. Following this, samples were normalized to 5 ng/µL for library prep. Following quantification, 30 samples were processed further. Libraries were prepared using a modified version of the Nextera protocol from Illumina (Therkildsen and Palumbi, 2017) (Supplementary Materials). Libraries were shipped to the Genewiz facility in South Plainfield, NJ, USA for sequencing on a NovaSeq 6000 S2 platform using 2 x 150 chemistry.

For samples from 2019, we performed phenol-chloroform DNA extractions (Sambrook and Russell 2009). DNA quality was assessed via Thermo Scientific Nanodrop Eight Spectrophotometer and DNA concentration was determined using an Invitrogen Qubit 4 Fluorometer. Purified DNA was then sent for library preparation and sequencing on an Illumina NovaSeq (S4) at the UC Davis DNA Technologies and Expression Analysis Core.

To facilitate analyses requiring either an outgroup species or population (e.g., Population Branch Statistic, demographic analyses), we obtained data from 31 additional individuals from the Short Read Archive (SRA). These previously published data consisted of one *G. wheatlandi* male (Sardell et al., 2021) and 30 anadromous *G. aculeatus* from Baie-de-L’Isle-Verte, Quebec, Canada (Sylvestre at al., 2023). All samples used in this study and their locations are shown in Figure 1 with additional details described in Table S1.

**Figure 1.**
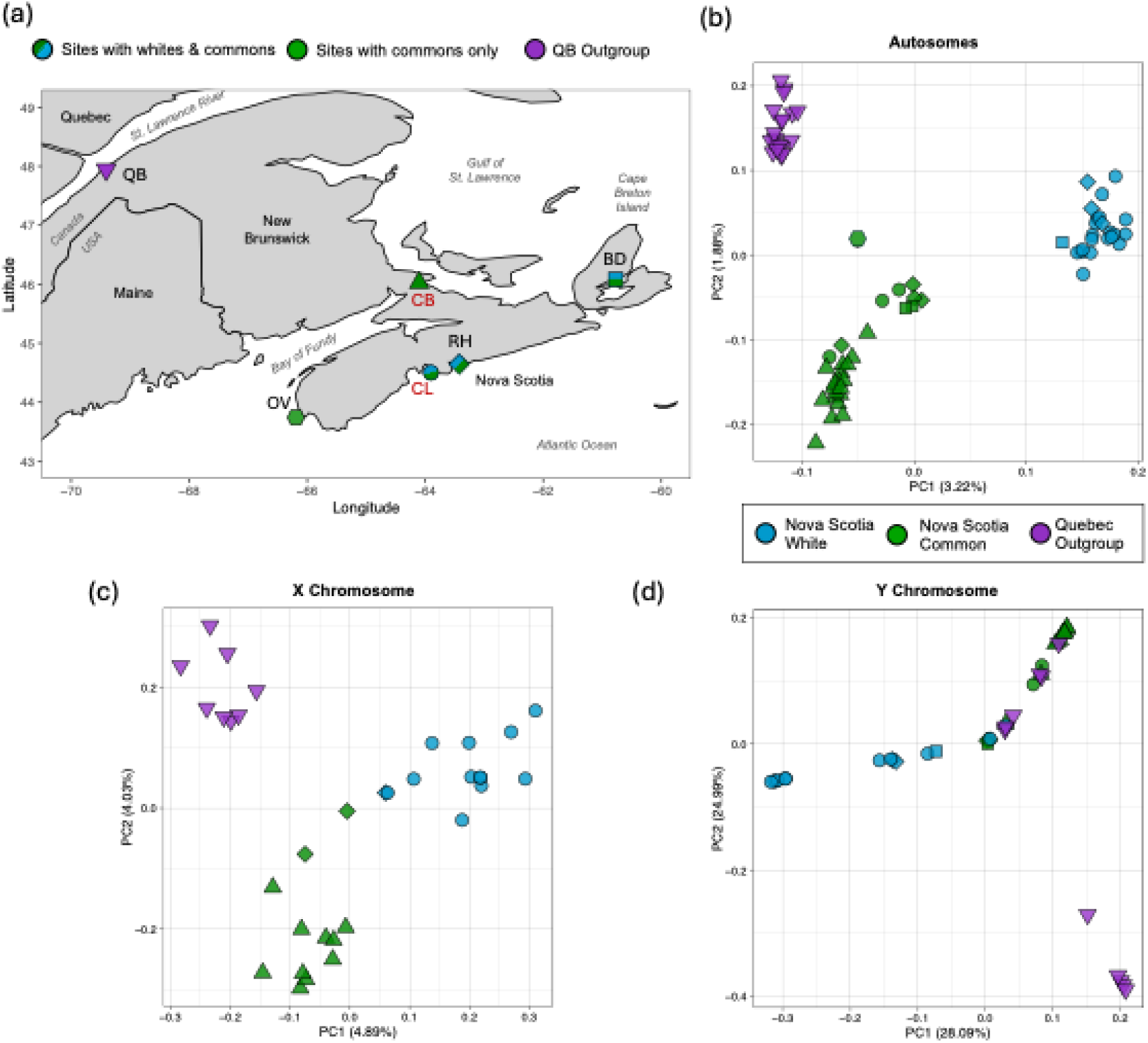
(a) Sampling sites of stickleback populations used in this study. Site labels: CB = Cherry Burton Road (triangle), OV = Overton (hexagon), CL = Canal Lake (circle), RH = Rainbow Haven (diamond), BD = Baddeck (square), QB = Baie-de-l’Isle-Verte (upside-down triangle). CB and CL are bolded in red as the majority of analyses were performed on these individuals. (b – d) Principal Component Analyses of the autosomes, and X and Y chromosomes of white and common sticklebacks from various locations in Atlantic Canada. (b) PCA of 1,067,766 autosomal SNPs from 19 Quebec outgroup (purple), 29 Nova Scotia common (green), and 23 Nova Scotia white sticklebacks (blue). (c) PCA of 41,635 SNPs from the X chromosome of 12 Nova Scotia common, 13 Nova Scotia white, and eight Quebec outgroup female sticklebacks. (d) PCA of 5,232 SNPS from the Y chromosome of 16 Nova Scotia common, 14 Nova Scotia white, and 11 Quebec outgroup male sticklebacks.

### Variant identification and processing

Male and female sequence reads were separately aligned to sex-specific v5 stickleback references (Nath, et al., 2020;1 Peichel et al., 2020) to correct for bias in mapping to the sex chromosomes (Webster et al., 2019). The modified sex-specific references were made by masking the Y chr with Ns for the female reference and masking the pseudoautosomal region (PAR) in the Y chr with Ns for the male reference based on coordinates from Peichel et al. (2020). Reference mapping for each sex was carried out using bwa mem (Li, 2013) and duplicate reads were removed using Picard (Broad Institute, 2019). Variant calling was performed using GATK HaplotypeCaller (Poplin et al., 2018). All prior steps were carried out using the snakemake pipeline, grenepipe (Czech and Exposito-Alonso, 2022). The resulting GVCFs for both sexes were then merged using the GenomicsDBImport argument in GATK. This was done for each chromosome prior to joint genotyping using GenotypeGVCFs. We specified the all-sites vcf option to ensure the final VCFs included both variant and invariant sites.

The resulting variant site VCFs were then subset and filtered in VCFtools (Danecek et al., 2011) depending on the analyses they were used for. In general, we retained variants that were biallelic SNPs and followed the GATK best practices hard filters (Geraldine et al., 2018). For demographic analyses exclusively, we did not apply a minor allele frequency (MAF) filter. Unless otherwise stated, the sex chromosomes were excluded and filtered separately from the autosomes. Analyses on the X chromosome (chromosome XIX in threespine stickleback) only utilized female samples, and analyses on the Y chromosome only utilized male samples. For all analyses on the Y chromosome, heterozygous calls were removed, and all sites were converted to haploid calls using BCFtools (Danecek et al., 2021) +fixploidy. The final dataset consisted of 1,067,766 SNPs for analyses on the autosomes, 41,635 SNPs for the X chromosome, and 5,232 SNPs for the Y chromosome. All variant calling and filtering for SNPs was performed on the University of California, Riverside High-Performance Computing Cluster.

### Population Genomics

To confirm if whites and commons represent distinct genetic clusters as in Samuk (2016), we performed principal component analysis (PCA) using the program snpRelate (Zheng et al., 2012) in R. PCA was conducted on SNP datasets of the autosomes and sex chromosomes for three populations (Quebec, Nova Scotia Commons, and Nova Scotia Whites), and then for 40 individuals (20 male and 20 females) from Canal Lake (CL) and Cherry Burton Road (CB) (Figure 1, S1).

We then calculated Weir and Cockerham’s F_ST_ (Weir and Cockerham, 1984) in 50kbp windows for the autosomes and sex chromosomes of whites and commons from CB and CL using pixy (Kournes and Samuk, 2021). All negative F_ST_ values for this analysis and those following were set to 0. F_ST_ is a relative metric of differentiation and high values can result from low within population nucleotide diversity (π; Nei and Li, 1979) caused by low recombination, local adaptation, and other evolutionary processes (Cruickshank and Hahn, 2014; Noor and Bennet, 2009). Thus, we also calculated π and *d_XY_* (Nei and Li, 1979)) for both ecotypes. In order to use pixy, we filtered invariant and variant site VCFs separately and then combined them using the BCFtools concat function. Nucleotide diversity in the PAR of the X chromosome of humans has been shown to be significantly higher than in regions of the X that do not recombine with the Y (Cotter et al., 2016). To account for this bias in analysis of the X chromosome, we calculated π in the PAR, which spans approximately 2.5 Mb (Roesti, et al., 2013), separately from the rest of the X chromosome excluding the PAR (nonPAR).

### Demographic Model Fitting with dadi

To estimate rates of gene flow and the timing of divergence between whites and commons, we fit a series of demographic models using the command line interface of the program dadi (Gutenkunst et al., 2009). We used a dataset composed of the autosomes from 40 white and common individuals from CL and CB, and one *G. wheatlandi* as an outgroup. We used SnpEff (Cingolani et al., 2012a) to predict the functional effect of each variant and used SnpSift (Cingolani et al., 2012b) to generate a VCF for dadi with only synonymous sites, which were assumed to be evolving neutrally or nearly neutrally. This annotated VCF was then converted back to the genome coordinates of the stickleback v5 assembly using CrossMap (Zhao et al., 2013). Finally, we polarized this VCF using Maximum Likelihood Ancestral Annotation in fastDFE (Sendrowski and Bataillon, 2024). We specified *G. wheatlandi* as an outgroup population and used the “n_target_sites” argument to ensure enough monomorphic sites were included in the analysis. This final polarized VCF was used to generate a 2D unfolded site frequency spectrum (SFS) in dadi-cli.

Using dadi-cli, we then inferred the fit of 11 demographic models of varying complexity to the unfolded spectrum. Inference for each model was run for 200 optimizations and then assessed for convergence. We specified the “--global-optimization” option to increase the chances a model converged and compared the fit of each model to our data using AIC. For the top three models, confidence intervals for each parameter estimate were inferred by generating bootstrap replicates and using the Godambe Information Matrix (GIM) in dadi-cli.

### Population Size Reconstruction

To further explore the demographic histories of whites and commons, we reconstructed changes in their independent effective population sizes over time using the pairwise Markovian sequentially coalescent (PSMC; Li and Durbin, 2011). PSMC relies on the coalescent to estimate historical demography from a single individual and has been shown to infer more ancient demographic events with greater accuracy compared to those that rely exclusively on the SFS (Mather et al., 2019). Thus, we chose this method as a complement to the dadi analysis.

PSMC input files were generated from four individuals total, one male and one female common, and one male and one female white. Each individual bam file was first filtered to include only the autosomes. We then used BCFtools mpileup and call to generate a VCF. The VCF were then converted to a diploid fastq file using the vcf2fq function. We specified the same average depth of coverage using ‘-D = 72’ and ‘-d = 30’ that was used when filtering datasets used for other analyses. PSMC was then run with default settings with 30 rounds of bootstrapping. PSMC curves were scaled and plotted using the same mutation rate and generation time as in the dadi analysis.

### Genome Scans for Candidate RI Loci

To identify regions of the genome potentially involved in RI between whites and commons (i.e., outlier loci), we calculated genome-wide Population Branch Statistic (PBS; Yi et al., 2010). Compared to pairwise metrics of divergence (i.e., F_ST_), PBS leverages an “outgroup” population to examine lineage-specific differences and has been shown to be a powerful method for detecting recent selection (Yi et al., 2010). We used a dataset composed of sticklebacks from Quebec as an outgroup population, and whites and commons sampled from multiple locations in Nova Scotia, Canada. Because this dataset included sticklebacks sequenced at different coverages, we performed additional SNP filtering in the R program SnpfiltR (DeRaad, 2022). Individuals with a proportion of missing data over 0.1 were removed from the analysis and we applied a missing data cutoff of 0.95. We performed a PCA using snpRelate to check that populations of each ecotype clustered together prior to calculating PBS. Next, we took the following steps to calculate PBS for whites: (1) We calculated Weir and Cockerham’s F_ST_ between the Quebec outgroup, Nova Scotia whites, and Nova Scotia commons (three pairs total) using VCFtools. (2) We transformed these pairwise F_ST_ estimates for each population pair into branch lengths (T), using the formula: T = − *log* log(1 − *Fst*). (3) PBS for white stickleback was then computed using the formula:

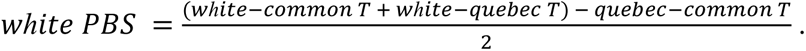

All negative PBS values were set to 0. We repeated the procedures above to calculate PBS for the sex chromosomes. As a comparison to the lineage specific differences in whites, we also calculated genome-wide PBS for Nova Scotia commons and the Quebec outgroup.

PBS values above the 0.99 quantile were considered outliers. To assess whether PBS outliers for whites overlapped with genes involved in phenotypic divergence and prezygotic isolation, we used a custom R script to identify genes and their Gene Ontology (GO) terms using the R package biomaRt (Durinck et al., 2009). We then searched for enriched GO terms among PBS outliers using g:-Profiler (Raudvere et al., 2019) and corrected for false positives using the g:SCS algorithm. P-values below the 0.05 threshold were considered significant. Lastly, we used SnpEff to predict the functional effect and chromosomal position of each PBS outlier. Because SnpEff uses a prior assembly version of the threespine stickleback, we used CrossMap to convert genome coordinates from the v5 assembly to the BROADS1 assembly.

### CNV Calling and Population Genomic Analyses

In addition to the SNP analyses, we identified copy number variants (CNVs) and inversions in our highest coverage dataset of whites from CL (n = 20) and commons from CB (n = 20). CNVs and inversions were called and genotyped with default settings in Smoove v0.2.8 (Pedersen et al., 2020). We generated separate VCF files for the autosomes with all individuals, for the X chromosomes using only females, and for the Y chromosome using only males. Using VCFtools v0.1.14, we filtered resulting VCF files by variant quality (--minQ 1) and required the ALT allele to occur at least eight times across all individuals (--non-ref-ac 8).

Relative differentiation in CNV allele frequencies between commons and whites was calculated across the autosomes and sex chromosomes separately using Weir and Cockerham’s F_ST_ in VCFtools. PCA of deletions and duplications on the autosomes and sex chromosomes were generated using Plink v1.9. F_ST_ values above the 0.99 quantile were considered CNV outliers. Interestingly, we encountered six CNV outliers greater than 100 kb including three approximately 3–7 mega base pairs long deletions on chromosomes I and IV. To account for the possibility these calls were false positives, we cross validated the CNV regions called from Smoove with another CNV caller, CNVpytor v1.3.1 (Suvakov et al., 2021). We considered calls with > 50% CNV length overlap between Smoove and CNVpytor with both males and females to be high-confidence CNV calls. As a complement to allele frequency differentiation of CNVs, we measured ecotype differentiation in copy numbers using a custom R script to calculate V_ST_ at CNV loci (Redon et al., 2006). Lastly, following the same procedure as in the SNP analyses, we identified genes overlapping or near CNV outliers and retrieved GO terms.

## RESULTS

### Whites and commons form distinct genetic clusters

PCA revealed that whites and commons form distinct genetic clusters irrespective of their geographic location (Figure 1b-d, S1). The inclusion of the Quebec outgroup resulted in three discrete clusters with PC1 explaining the divergence between Nova Scotia commons and Nova Scotia whites. This pattern was most clear on the autosomes with PC1 explaining 3.2% of the variation (Figure 1B). Population membership on the X chromosome, which included females exclusively, exhibited moderately discrete clustering with the Quebec outgroup forming the most distinct cluster. The Y chromosome, which included males exclusively, exhibited groupings that were less clear with several Quebec outgroup individuals and one Nova Scotia white grouping with Nova Scotia commons. On the X and Y chromosomes, PC1 explained 4.89% and 28.09% of the variation respectively (Figure 1c and d). In these PCAs of the sex chromosomes, many individuals grouped towards zero likely due to missing data.

Similarly, in the PCA of the autosomes of only the CL (white) and CB (common) populations, whites and commons formed clearly distinct clusters with PC1 explaining 4.89% of the variation (Figure S1a-c). This pattern was reflected on the X chromosome of CL and CB females with PC1 explaining 6.94% of the variation (Figure S1b). On the Y chromosome, there were two distinct clusters of whites and commons separating on PC1 and a third diffuse group of several whites and one common that separated on PC2 (Figure S1c).

### White and commons diverged recently under an isolation-migration (IM) model

The model fitting approach suggests that an IM model best explains the nature of divergence between whites and commons and that divergence occurred approximately 623 ya but could have occurred anytime from now to ∼1,277 ya (Figure 2, Table 2 and S2). The IM model detected low levels of gene flow from whites to commons (∼ 0.0017 migrants per generation), but even lower levels of gene flow from commons to whites (∼ 0.0005 migration per generation). N_e_ of whites was over twice as high than commons, but both experienced large population expansion since divergence. The second-best model was a strict isolation model where speciation occurred approximately 421 ya (∼383 – 458.87). Similar to the IM model, white N_e_ was much larger than commons. The third-best model was an ancient symmetric migration model where migration occurred once ∼350 ya and again ∼66 ya respectively. The number of migrants per generation was extremely low and white N_e_ was nearly twice as high as commons.

**Figure 2.**
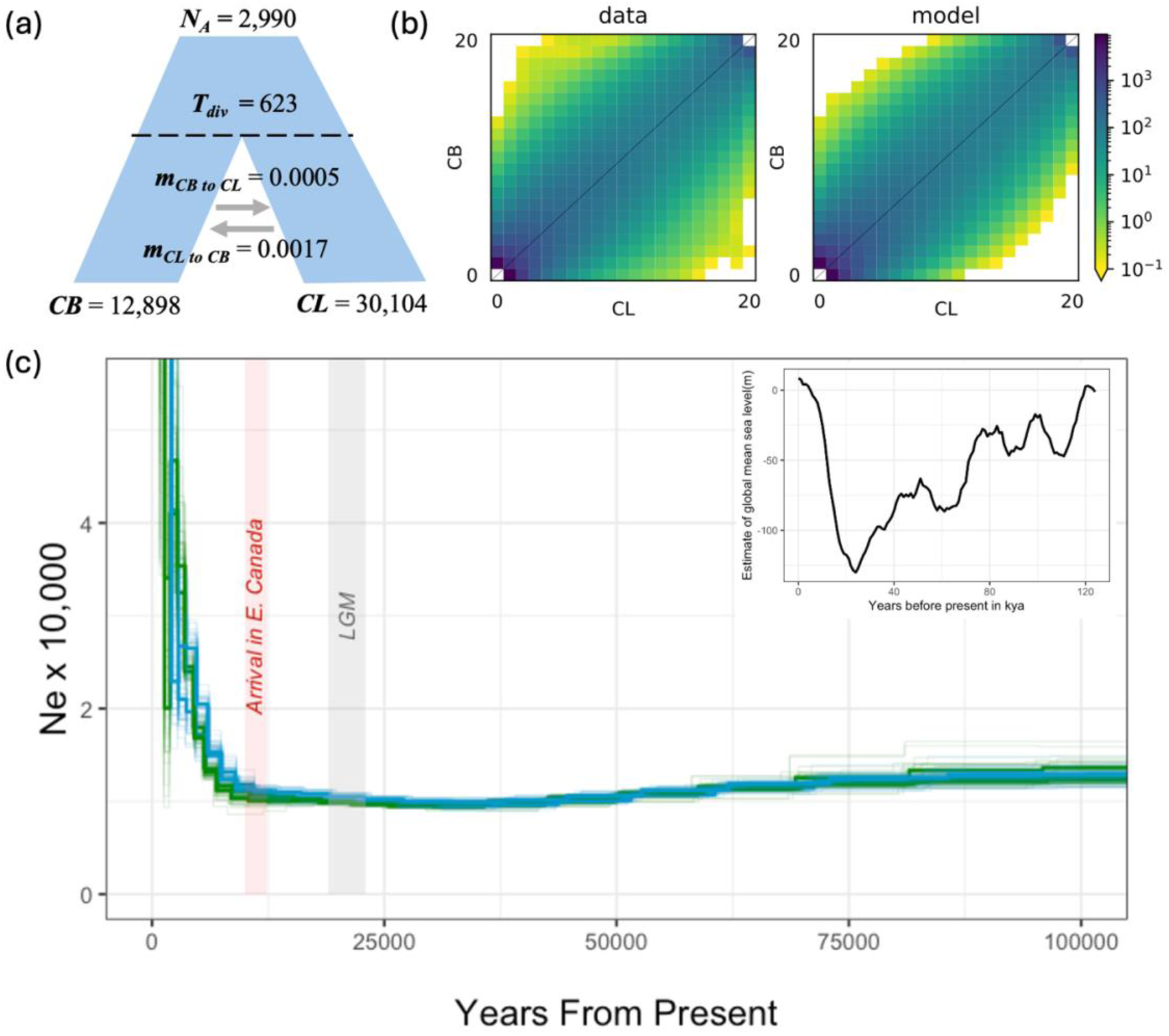
Demographic analyses estimating the nature of divergence and effective population size histories of white and common stickleback. (a) Schematic of isolation migration model determined to be the most likely scenario of divergence between white and common stickleback by AIC with scaled, mean values. (b) the observed (left) and simulated (right) polarized 2D site frequency spectrum for white (CL on x-axis) and common (CB on y-axis) sticklebacks inferred using dadi -cli. The joint allele frequency in each 2D bin is indicated via color scale (darker = higher frequency). (c) Effective population size histories for two common (green lines) and two white (blue lines) sticklebacks estimated in PSMC. Thinner lines represent bootstrap replicates. Years ago scaled by generation time and mutation rate are on the x-axis, and N_e_ is on the y-axis. Grey shading represents the estimated range of the Last Glacial Maximum (19 – 23kya) and red shading represents the time period (∼10–12kya) when stickleback became established in eastern Canada (Haines, 2023). The smaller box in the top right corner represents global mean sea level estimates (Spratt and Lisiecki, 2016).

**Table 1.** Genome-wide averages of F_ST,_ *d*_*XY*,_ and π describing divergence between white and Nova Scotia common sticklebacks in the autosomes, X chromosome, and Y chromosome. Mean and the upper and lower limits of the 95% confidence interval are shown for each summary statistic. 95% confidence intervals were calculated from 10,000 bootstrap replicates.

| Autosomes (male and female) |  |  |  |  |  |
| --- | --- | --- | --- | --- | --- |
| $F_{ST}$ | $d_{XY}$ | White $\pi$ | | Common $\pi$ | |
| 0.0343 ± | 0.037 ± | 0.0358 ± | | 0.0363 ± | |
| 0.0011 | 0.00045 | 0.00045 |  | 0.00045 |  |
| X Chromosome (female only) |  |  |  |  |  |
| F<sub>ST</sub> | $d_{XY}$ | White $\pi$<br>nonPAR | White $\pi$ PAR | Common $\pi$<br>nonPAR | Common $\pi$<br>PAR |
| 0.0263 ±<br>0.0035 | 0.0281 ±<br>0.0021 | 0.0208 ±<br>0.00105 | 0.0731 ±<br>0.0067 | 0.0211 ±<br>0.00105 | 0.0735 ±<br>0.00665 |
| **Y Chromosome (males only)** |  |  |  |  |  |
| F<sub>ST</sub> | *d<sub>XY</sub>* | White $\pi$ | Common $\pi$ | | |
| 0.2529 ±<br>0.0136 | 0.013 ±<br>0.0012 | 0.0106 ±<br>0.00115 | 0.006 ±<br>0.0003 |  |  |

**Table 2.** Demographic model parameters and scaled parameters estimated by dadi-cli for the top three demographic models. Values in each cell represent scaled parameter values with respect to commons (population 1) and whites (population 2). θ, 4N_e_u; N_anc_, effective population size of the ancestral population; s, relative size of population 1 after split (size of population 2 1-s); nu1 and nu2 size of population 1 and 2 at present; m, migration (individuals per generation) between pop1 and 2, m_12_, migration rate from population 1 to population 2; m_21_, migration rate from population 2 to population ; T divergence time (generations) between population 1 & 2, T1, time between divergence and secondary contact; T2 time between secondary contact and present. Log likelihoods (LL) and Akaike Information Criterion (AIC) scores are shown for each model.

| Parameters | IM | Strict Isolation | Ancient Symmetric Migration |
| --- | --- | --- | --- |
| $N_{anc}$ | 2990.17 | 3013.28 | 3015.366 |
| $s$ | 1001.68 | — | — |
| $nu1$ | 12897.73 | 9544.083 | 10166.65 |
| $nu2$ | 30104.16 | 43189.41 | 29664.19 |
| $m$ | — | — | 0.00004 |
| $m_{12}$ | 0.0017 | — | — |
| $m_{21}$ | 0.0005 | — | — |
| $T$ | 622.83 | 417 | — |
| $T1$ | — | — | 66.45 |
| $T2$ | — | — | 354 |
| $misid$ | 0.043 | 0.043 | 0.043 |
| $\theta$ | 8683.75 | 8750.87 | 8756.92 |
| **Log Likelihood** | -1193.96 | -1225.36 | -1225.2 |
| **AIC** | 2401.92 | 2458.71 | 2462.39 |

Whites and commons appear to have nearly identical N_e_ histories based on the PSMC graphs except for within the last 10 kya where there is more uncertainty (Figure 2). Fang et al., (2018) estimated Western Atlantic stickleback (Long Pond, Nova Scotia, Canada; the St. Lawrence River, Quebec, Canada; and Penobscot Bay, Maine, U.S.A.) and populations from the Barents and Norwegian Seas diverged between 7.1 and 17.1 kya. Based on this phylogenetic evidence and fossil stickleback from near the Ottawa River dated to ∼10 kya (Dyke 2004; McAllister et al., 1981), Haines (2023) proposed that stickleback became established in eastern Canada 10-12.4 kya. The PSMC graphs show that both populations experienced a very slight bottleneck starting approximately 60 kya followed by a drastic population expansion approximately 20 kya (Figure 2c). The timing of this population growth coincides with the end of the Last Glacial Maximum (LGM) and dates suggested by Haines (2023).

### Genome scans exhibit small, localized peaks amid low genetic divergence

The F_ST_ genome scan between CL and CB populations revealed seven to 10 small regions across the genome with near fixed differences including on the Y chromosome (Figure 3, top panel). This stood in contrast to our expectation that we would find high F_ST_ exclusively in one to three localized regions of the genome such as within chromosomal inversions. Mean F_ST_ was extremely low (<0.05) on the autosomes and X chromosome, but much higher on the Y chromosome (∼0.25). Mean *d_XY_* (0.037) and π of whites (0.0358) and commons (0.0363) was similarly low on autosomes (Table 1). On the X chromosome, mean white π (0.0208) was slightly lower than common π (0.0211) in nonPAR regions. For both populations, π in the PAR (∼0.073) was over three times higher than in the nonPAR region (∼0.021). Mean *d_XY_* on the Y chromosomes (0.013) was lower than on both the autosomes (0.037) and X chromosomes (0.028), and π of male whites (0.011) was nearly 2 times higher than π in male commons (0.006) on the Y chromosome.

**Figure 3.**
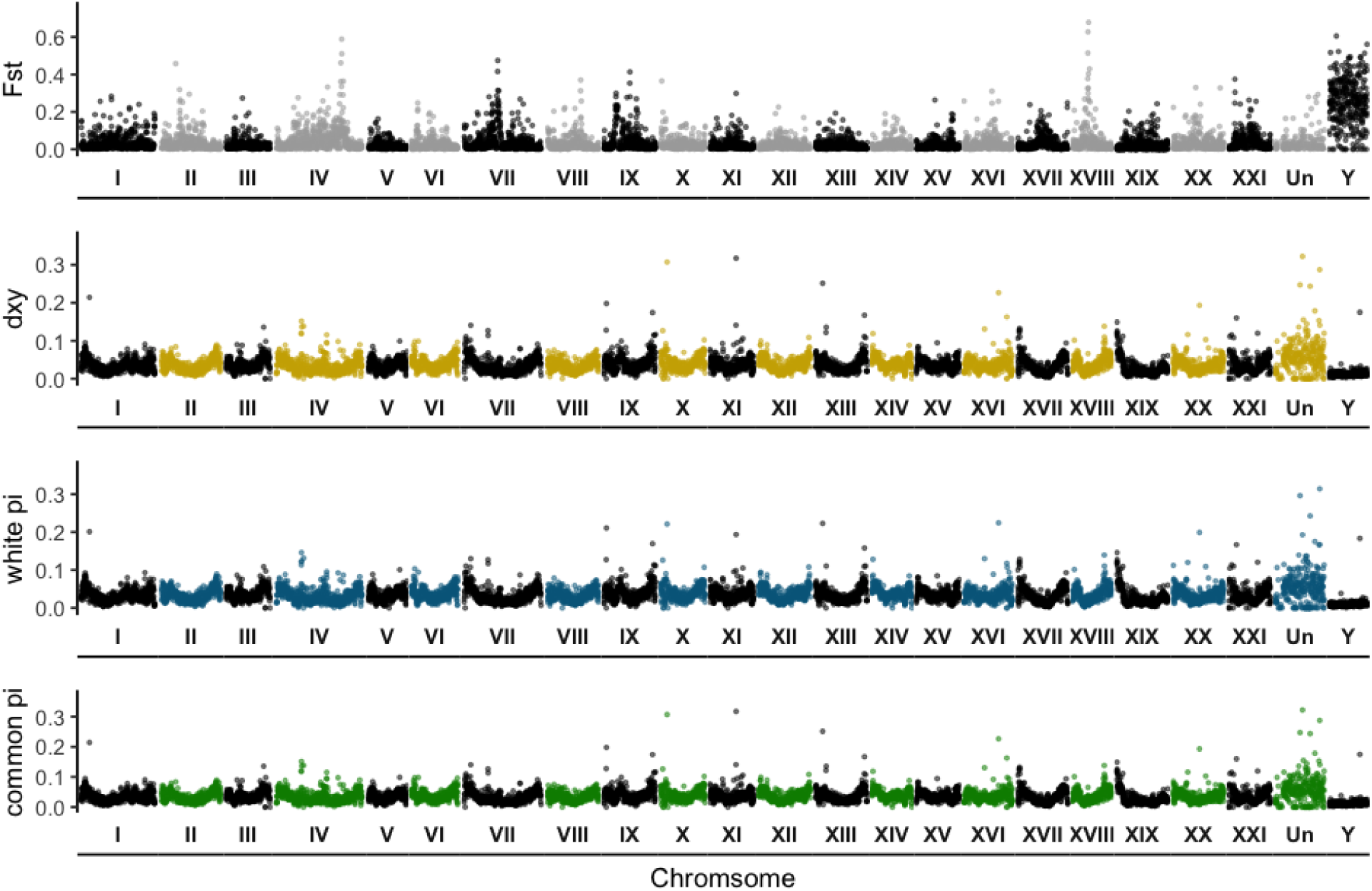
Pairwise summary statistics describing genome-wide divergence between white and common sticklebacks from CL and CB respectively. F_ST,_ *d*_*XY*,_ and π in white and common sticklebacks are distinguished by different colored points. Summary statistic values are plotted in genomic order within each chromosome (in Roman numerals and demarcated by space), with “Un” representing concatenated unplaced scaffolds. Divergence metrics were calculated in 50k bp windows for the autosomes, the X chromosome (XIX, which includes PAR and nonPAR), and the Y chromosome. The X and Y chromosomes were composed of only female and only male individuals respectively and were filtered and processed separately before plotting together with the autosomes.

Similar to the F_ST_ scan, there were ten to fifteen small regions of high PBS across the genome for whites (Figure 4, Table S3). Average PBS of whites (0.0263) on the autosomes was higher than the Quebec outgroup (0.0176) and over twice as high as Nova Scotia commons (0.0102). On the X chromosome, average PBS was similar between the three populations (0.02-0.03). On the Y chromosome, average PBS was similar between whites (0.1772) and Nova Scotia commons (0.1602), but approximately 5 times lower than the other populations for the Quebec outgroup (0.0317). F_ST_ between the three population pairs (Nova Scotia commons, Nova Scotia whites, and Quebec) used to calculate PBS (Table S4) was generally very low across autosomes and the X (<0.05), especially between the Quebec outgroup and Nova Scotia commons on the autosomes (<0.02). F_ST_ was highest for all population pairs on the Y chromosome (>0.1).

**Figure 4.**
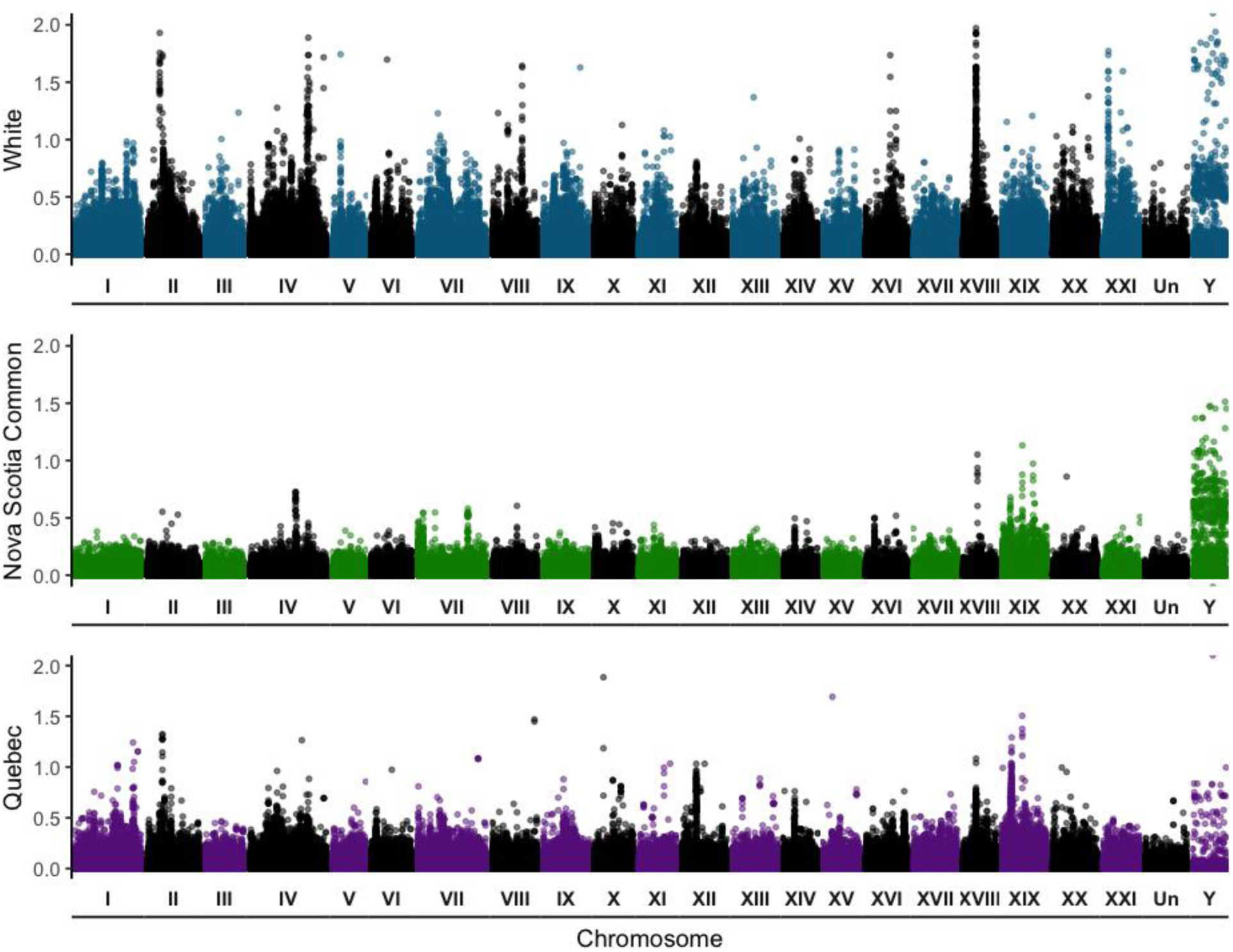
Population branch statistic (PBS) estimated for the autosomes and sex chromosomes of Nova Scotia white, Nova Scotia common, and Quebec outgroup sticklebacks. The autosomes and sex chromosomes (XIX and Y) were filtered separately before calculating PBS and plotting them together.

### Patterns of differentiation within CNVs corroborate SNPs

Among 40 individuals from CL and CB, we identified 94,754 deletions and duplications and 1,724 putative inversions. Unlike the deletions and duplications, the majority of inversions exhibited F_ST_ values near 0 with the highest value being 0.37. Because the putative inversions were less differentiated and we lacked the long read sequencing data to validate them, we focused all downstream analyses on deletions and duplications.

Similar as with SNPs, PCA of the CNV autosomes revealed that whites and commons formed distinct clusters, with PC1 explaining 6.95% of the variation (Figure 5a). This pattern was also seen in the PCA of the X chromosome (Figure 5b). PCA on the Y chromosome revealed less distinct clustering with two whites grouping near commons and one common separating from both white and common clusters (Figure 5c). Mean CNV F_ST_ and V_ST_ were extremely low (<0.1) for the autosomes, X chromosome, and Y chromosome (Table S5). Similar to the SNP data, F_ST_ and V_ST_ were highest on the Y chromosome and lowest on the autosomes. F_ST_ and V_ST_ were positively correlated (Figure S2a). Genome-wide F_ST_ of the CNVs revealed a similar genomic landscape to both the F_ST_ and PBS analyses of the SNPs where differentiation was high in several locations across the genome rather than in a few localized regions (Figure 5d).

**Figure 5.**
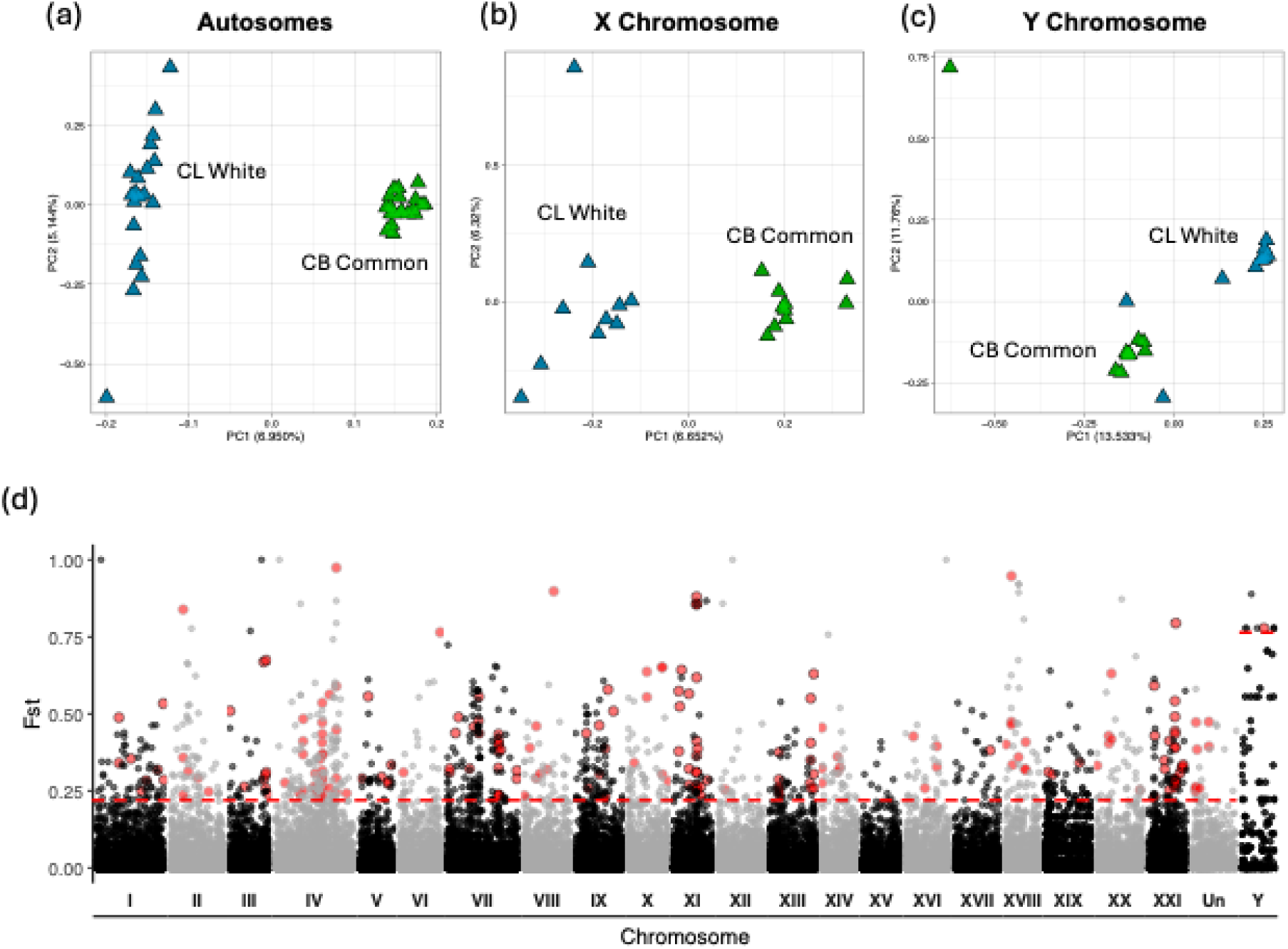
Genomic divergence in deletion and duplications between 20 white and 20 common sticklebacks from CL and CB respectively. PCA of CNVs on the (a) autosomes, (b) the X chromosome of females only, and (c) the Y chromosome. (d) Genome-wide F_st_ for deletions and duplications. Red dotted lines represent the 0.99 quantile of PBS values with values above the threshold being considered outliers. Larger red dots represent CNV outliers that have been cross-validated.

The CNV outlier analysis identified 117 duplications and 836 deletions across the autosomes and sex chromosomes combined. After cross-validation across CNV variant callers, 74/117 duplications and 132/836 deletions remained, with nine deletions on the Y chromosome and 20 deletions and two duplications on the X chromosome. Outlier duplications ranged from 132,461bp spanning six genes to 285 bp with mean length of 17,383bp. Deletions ranged from 75,223bp spanning three genes to 31 bp long with mean length 3,214 bp (Figure S2a). The mean distance of CNVs from genes was 12,117bp, and 63 CNVs overlapped with genes (Figure S2b).

### Gene function analysis suggests processes involved in RI

A total of 1,044 genes overlapped with white PBS outliers on the autosomes (Table S6). We found genes with major functions in reproduction such as androgen binding, gonad development, spermatogenesis, estradiol binding, and estrogen signaling. Several candidate genes were also linked to sexual signaling and sensory modalities including melanin pigmentation, olfaction, photoreception, hearing, and brain development. On the X chromosome, 78 genes corresponded to PBS outliers, which included one gene involved in egg coat formation and sperm-egg binding, a heat shock protein (hsf4) involved in lens development, and a gene (galn) involved in galanin signaling. Eight genes, of which three had annotated functions (hepatocyte growth factor (HFG), protein phosphatase 6 regulatory subunit 2 (ppp6r2b), and SLIT-ROBO Rho GTPase activating protein 1a (srgap1a) corresponded to PBS outliers on the Y chromosome.

The GO enrichment analysis revealed that white PBS outliers were enriched for “calcium ion binding”, “signaling receptor activity”, “homophilic cell adhesion via plasma membrane adhesion molecules”, “signaling”, “response to stimulus”, and “membrane” (Table S6). Based on the SNPeff annotations of the PBS outliers, only 2.58% of variants were in coding regions with the remaining variants being in non-coding regions such as intergenic regions (41.23%) and introns (26.35%). This suggests that these mutations mainly act on gene regulation and expression.

Several CNV outliers overlapped genes with relevant functions to RI such as egg coat development and sperm binding, olfaction, eye development, sound perception, and swimming behavior (Table S7). The GO enrichment analysis revealed that CNVs were enriched for “sialytransferase activity”, “olfactory receptor activity”, “toxin activity”, “detection of chemical stimulus involved in sensory perception”, “modulation of process of another organism”, and “membrane” (Table S7). While it is less clear how the other GO terms relate to RI in this system, “olfactory receptor activity” and “detection of chemical stimulus involved in sensory perception” can be clearly linked to potential prezygotic barriers.

## Discussion

Incipient species offer the unique opportunity to explore the initial genetic changes that underlie RI. We leverage the young white stickleback system from Nova Scotia, Canada to explore patterns of genomic divergence and use demographic modeling to infer their evolutionary history. Our results align with prior genomic studies that demonstrate the distinctness of whites and commons, and suggest these lineages diverged very recently (< 1k ya). Genome-wide F_ST_ is extremely low except for in a few small peaks spread across the genome. A novel finding from our study is that CNVs and SNPs within this system exhibit similar patterns, which bolsters evidence for the distinctness of these ecotypes. Furthermore, we identified outliers with functions that could be important for RI between ecotypes. Unlike most population genomic studies of sticklebacks, we incorporated the Y chromosome in our analyses and found less discrete population membership. We discuss the significance of these findings and highlight unexplained patterns and future areas of research below.

### The recent postglacial origin of white and commons and their phylogeographic structure

A major objective of our study was to confirm that the white stickleback represents an incipient species by estimating the timing and nature of their divergence. Unsurprisingly for recently diverged populations, PSMC revealed nearly identical demographic histories of whites and commons, especially during more ancient times. Very recent divergence times (∼1,277 to now) were estimated in dadi. Similar to Samuk (2016), we found evidence for speciation with gene flow although our estimates of the number of migrants per generation were much lower. This system adds to a growing number of young, sympatric stickleback ecotypes that are not ecologically distinct. For example, stickleback ecotypes from the manmade Jordeweiher Pond in Switzerland (Feller et al., 2017; Marques et al., 2016) diverged ∼100 ya. These ecotypes exhibit both divergent nuptial coloration and nesting behaviors but lack ecological differences.

Our results support an evolutionary scenario where the ancestors of whites and commons colonized the western Atlantic from northern Europe following deglaciation of the St. Lawrence valley approximately 10-12 kya (Haines, 2023). During this time, they experienced a significant population expansion where they spread along the Atlantic coast into what is now their contemporary range. This pattern of rapid postglacial expansion has been observed in other PSMC analyses on the whole genomes of marine stickleback from Greenland and Northern Europe (Liu et al., 2016). It is also reflected in the dadi results where modern whites and commons have much higher N_e_ compared to their ancestors.

A caveat of most demographic analyses is that inference can be impacted by natural selection, population structure, and gene flow (reviewed in Beichman et al., 2018). To account for biases caused by temporal and genomic changes in N_e_ and migration rate, we attempted to incorporate more “realistic” models into our dadi analysis (Rougeux et al., 2017, Momigliano et al., 2021). Unfortunately, these failed to converge. Confidence intervals inferred by GIM for the IM model are very large compared to the other two best-fitting models. Regardless, the AIC scores between these three models are very similar and estimate similar patterns of low gene flow and higher N_e_ in whites compared to commons. Clearly, the demographic history of this system is very complex due to its recent divergence, high historical levels of gene flow, and recent population expansion that is characteristic of marine stickleback. These complexities are only partially captured in our analyses.

Lastly, our results add to several studies demonstrating that commons from Nova Scotia and populations from the western Atlantic represent multiple phylogeographic lineages rather than a single panmictic population (Behrens et al, in review 2025b; Fang et al., 2018; Haines, 2023; Samuk, 2016). Individuals from the Bras d’Or Lake, an inland sea on Cape Breton Island, form separate genetic clusters from commons from mainland Nova Scotia (Behrens et al., in review 2025b; Samuk, 2016). Similar to the phylogenetic analysis of Fang et al. (2018), the Quebec outgroup was distinct from Nova Scotia commons. Based on two freshwater (Long Pond) and three marine individuals (Penobscot Bay and the St. Lawrence River), Fang et al. (2018) suggested that western Atlantic stickleback populations diverged between 4.5 and 12.4 kya. Pairwise F_ST_ between the Quebec outgroup and Nova Scotia commons is lower (∼0.016) than F_ST_ between Nova Scotia commons and whites (∼0.024). This suggests more consistent gene flow between these two populations or that they are more closely related despite being more geographically distant. These interesting patterns call for future research with more comprehensive sampling into the phylogeographic history of common populations in Atlantic Canada and surrounding regions.

### Concordant patterns of divergence in SNPs and CNVs are genome-wide

We predicted inversions would be important for facilitating RI in our systems and expected to see signatures of them in our genome scans (i.e., large “blocks” of F_ST_). Instead, we consistently recovered patterns of diffuse differentiation in many small peaks in both our F_ST_ and PBS scans. While we found putative inversions in our genomic data, we did not cross validate them with long read data. Relatively lower F_ST_ of these putative inversions compared to deletions and duplications suggest that the inversions were not highly differentiated between ecotypes. Together with the genome scans, it is unlikely that RI is facilitated by inversions in this system. Another prediction we had was that differentiation would be higher on the sex chromosomes. Instead, we found less differentiation on the X chromosome and differentiated genes relevant to prezygotic isolation (i.e., olfaction, vision) were spread across the genome rather than restricted to the sex chromosomes. This is similar to studies on *Drosophila* (Moehring et al., 2006) and *Lapulla* crickets (Shaw et al., 2007) where genes responsible for mating discrimination and preference are spread throughout the genome.

A potential explanation for these genome-wide vs more localized patterns is the genome-wide congealing (GWC) hypothesis (Feder et al., 2014). According to this hypothesis, the aggregate effects of many RI genes, rather than a few genes, can eventually lead to the genomes of speciating lineages “congealing” into distinct entities. GWC is thought to occur in two phases. In the “genic” phase, populations are partially isolated and exhibit heterogeneous patterns of differentiation in a few isolated regions, but overall background divergence is low. Populations in the genic stage tend to not cluster into distinct groupings. After passing a certain threshold, species enter the “genomic” phase with elevated F_ST_ and RI genes across the genome. The white stickleback system appears to be in the latter stages of the genic phase of GWC as most of the genome, except for in seven to 10 small regions, exhibits low F_ST_, but both populations consistently form discrete genetic clusters.

Lastly, there are relatively few studies comparing divergence between SNPs and CNVs, and evolutionary theory about the relative contribution of CNVs and SNPs in RI is lacking. Our population genomic analyses of deletions and duplications recapitulated patterns observed in the SNPs. While it is generally true that population structure in SNPs is reflected by CNVs (e.g., humans: Conrad et al., 2010; threespine sticklebacks: Chain et al., 2014; Chinese cattle: Zhang et al., 2020), these signals are not always congruent (e.g., American lobster: Dorant et al., 2020; fungi: Bazzicalupo et al., 2020; songbirds: Delmore et al., 2023). Congruent signals between SNPs and CNVs suggest there is strong divergent natural selection acting on loci involved in RI in this system. This is surprising given the very recent divergence between white and commons because they share a large amount of ancestral genetic variation. Our results serve as evidence that differences in CNVs may accumulate as quickly as SNPs and highlight the need for more empirical and theoretical studies comparing the relative roles of SNPs and CNVs in speciation.

### Divergence and persistence of the white stickleback: Possible mechanisms

Even though our study did not explicitly test the strength of prezygotic barriers or functionally validate candidate genes, there may be a strong sensory component to divergence in this system. Unlike the GO analysis of the outlier SNPs, outlier CNVs overlapped with or were located near genes enriched for olfactory receptor activity and the detection of chemical stimuli involved in sensory perception (i.e., olfaction and gustation). We identified both duplications and deletions differentiated between whites and commons spanning two novel olfactory genes (ENSGACG00000020542 and ENSGACG00000020543) on chromosome VII and one novel olfactory gene (ENSGACG00000006963) on chromosome XVI. Among its other functions, olfaction is involved in mate recognition and sexual selection in several systems including in sticklebacks (e.g., Rafferty and Boughman, 2006; Milinski et al., 2005).

Among our top five outlier CNVs, we identified a 35,617 bp duplication spanning two genes, adhesion G protein-coupled receptor L4 (adgrl4) and vitellogenin 3 (vtg3) on chromosome VIII. These genes were exclusively and always duplicated in whites compared to commons. Vitellogenins have been implicated in proper embryonic and larval development in European sea bass (*Dicentrarchus labrax*), zebrafish (*Danio regio*), Atlantic halibut (*Hippoglossus hippoglossus*), and European plaice (*Pleuronectes platessa*) (reviewed in Yilmaz et al., 2024). These results are interesting given the distinct differences in egg size and ovarian fluid between female whites and commons (Behrens et al., 2025a; Grant, 1993).

Previous studies on the integument of whites and commons revealed that male whites from Canal Lake, Rainbow Haven, and Baddeck had lower densities and reduced coverage of melanophores compared to sympatric commons (Haley et al., 2019). It is possible that the pearlescent nuptial coloration of male whites may have evolved under the sensory drive model where more conspicuous signaling traits are more likely to be favored because they are more easily detected by receivers and less likely to be obscured with distance or changes in habitat (Cummings and Endler, 2018; Endler 1992). Our genome scans of SNPs identified seven outlier genes with GO terms related to pigmentation including melanin concentrating hormone receptor 2 (mchr2) on chromosome IV. In addition to its role in pigmentation, melanin concentrating hormone has been strongly associated with motivational behaviors in animals including postpartum care (reviewed in Diniz and Bittencourt, 2017). Recently, Bell et al. (2025) detected higher levels of pituitary melanoconcentrating prohormone in male whites compared to male commons. We also identified the neuropeptide galanin (galn) as an outlier on the X chromosome. Galanin is an especially interesting candidate because it has been implicated in color pattern formation in zebrafish (Eskova et al., 2020), feeding and infanticide in rodents (Wu et al., 2014) and parental care in vertebrates (reviewed in Kaplan et al., 2025). Indeed, there is already evidence that galanin is important for initiating paternal behavior in commons, but not whites: male commons, but not male whites, exhibit increased activity of preoptic area galanin neurons post-spawning (Maciejewski et al., 2025).

Currently, the role of prezygotic barriers in maintaining white and common ecotypes remains inconclusive. However, the consistent genetic distinctness of ecotypes from multiple sympatric populations, rareness of hybrids in nature, and low levels of gene flow detected in our demographic analyses imply that extrinsic postzygotic barriers alone are likely not enough to maintain RI (Behrens et al. in review, 2025b). Given our results, we hypothesize that the impact of prezygotic barriers on RI in this system may be strongest in nature where environmental cues and ecologically relevant traits such as nesting microhabitat preference and use are involved (e.g., Bolnick et al., 2015; Dean et al., 2021). Taken together, natural history studies and/or more natural laboratory studies are needed to better understand the interplay between microhabitat use, sexual selection, and RI in this system.

### Unexplained patterns: Sex-ratio bias and the Y chromosome

While not the focus of our study, we calculated genetic diversity on the sex chromosomes and compared them to the autosomes to make predictions about the effective sex ratio of whites and commons. We expected whites to have a smaller N_e_ compared to commons given their more recent divergence. Given their conspicuousness, we also thought male whites would have lower N_e_ or a more female-biased sex ratio due to increased rates of predation. Instead, we found that the ratio of Y:A diversity in commons (0.16) was lower than both neutral expectations (¼; Webster and Wilson Sayres, 2016) and the ratio of Y:A diversity in whites (0.29). While not sex-specific, dadi models also consistently inferred that N_e_ in commons is lower than in whites.

One possible cause of these patterns is that the effective adult sex ratio of commons is more female-biased and there are fewer reproducing males. Lower overall N_e_ in commons could also be due to male commons spending more time out of the breeding pool due to paternal care, or it could suggest that commons from CB that we used in our demographic analyses represent a more isolated population. Male whites might exhibit a more male-biased sex ratio due to whites being able to reenter the mating pool shortly after courtship rather than providing parental care. Behrens et al. (2024) hypothesized that this could lead to greater male-male competition and stronger sexual selection in whites compared to commons. These findings agree with existing theory on the operational sex ratio (OSR; Emlen and Oring, 1997; Kokko and Jennions, 2008) that posits that having more reproductively available males in a population increases the intensity of male-male competition to attract mates.

While these inferred sex ratios are intriguing, more formal calculations with more male individuals are needed to determine if they are reliable. Currently, life history differences (e.g., sex specific mortality, lifespan, and sexual maturity) between whites and commons are largely unknown. Determining the extent of these differences would provide valuable insight into our results.

Finally, there was a consistent lack of clustering amongst ecotypes in all our PCAs of the Y chromosome with both SNPs and CNVs. One possible reason for this is that there are unsorted ancestral polymorphisms segregating in male sticklebacks from this region. Little is known about intraspecific variation in the Y chromosome of stickleback and most studies omit the Y chromosome from their analyses. Another possibility is that variant calling and genotyping on the Y chromosome resulted in inaccurate genotypes due to initially calling variants on the Y chromosomes as diploid and subsequently converting diploid calls to haploid. Overall, additional studies on the Y chromosome with more male individuals are needed to understand if these patterns are biological or an artifact of low sample sizes.

### Conclusion

Our study highlights the utility of the white stickleback system as a model for studying the genomics of RI in its early stages. This work represents the most recent and comprehensive population genomic study on this system to date. It is also one of few studies comparing the contributions of SNPs and CNVs to RI and examining intraspecific variation on the Y chromosome in sticklebacks. Furthermore, it adds to a small, but growing number of studies on the biology and evolutionary history of western Atlantic sticklebacks. Using a genomic approach, our study confirms both the distinctness of white from common sticklebacks and their recent divergence. We find evidence that these lineages speciated with gene flow, but that gene flow is currently very low. It is possible that environmentally mediated prezygotic barriers play a large role in preventing gene flow between these ecotypes. Ultimately, the unique natural history of this system enables a wealth of opportunities to understand processes adjacent to speciation including sexual conflict, sensory biology, and gene expression.

## Supporting information

Supplementary Materials (Low Coverage Library Preparation, Figures S1, S2, Tables S3-S5

## Acknowledgements

1. F. J. J. C. and M. J. O were supported by funding from NSF Grant 214459. Research reported in this publication was supported by NIGMS of the National Institutes of Health under award number 1R35GM139597 to AMB. We are grateful to Laura Weir, Anne Haley, Stanley King and Anne Dalziel for collecting samples for this study and providing information about collected individuals. Members of the Bentzen lab made the following contributions for pre-2019 samples: Ian Paterson isolated DNA and prepared sequencing libraries, Matt Penney shared protocols and Beth Watson shared data. Ellie Armstrong assisted with interpreting the PSMC analysis. Maria Akopyan, Mikhail Plaza, and members of the Samuk Lab all provided valuable comments to the manuscript.

Canal Lake individuals were collected under Department of Fisheries and Oceans Maritime permit #343930, Cherry Burton Rd. individuals were collected under Department of Fisheries and Oceans Gulf Region permit #SG-RHQ-19-008, and the Overton individual was collected under the animal ethic approval number 12-045 from Dalhousie University.

## Data Accessibility and Benefit-Sharing Statement

Raw WGS sequence data for samples collected are deposited in the NCBI Sequence Read Archive under BioProject: PRJNA1244922. Accession numbers of genomic samples generated from this project are listed in Table S1. Metadata and associated code can be found in the github repository: https://github.com/asumarli/White_Stickleback_Speciation_Genomics

## Author contributions

A.S., K. S, and A.M.B. conceptualized the project. C.B. performed field collections in 2019. G. L. performed DNA extractions for whole genome sequencing. A.S. processed the sequencing data for population genetic analysis. F.J.J.C. and M.J.O. performed CNV calling and calculated V_st_. A.S. performed analyses and data visualization. A.S. wrote the manuscript implementing comments and suggestions from all coauthors.

## References

Albert, A. Y. K., & Otto, S. P. (2005). Sexual selection can resolve sex-linked sexual antagonism. *Science (New York*, N.Y.), 310(5745), 119–121. 10.1126/science.1115328

Andrew, R. L., Kane, N. C., Baute, G. J., Grassa, C. J., & Rieseberg, L. H. (2013). Recent nonhybrid origin of sunflower ecotypes in a novel habitat. Molecular Ecology, 22(3), 799–813. 10.1111/mec.12038

Bazzicalupo, A. L., Thomas, M., Mason, R., Munro-Ehrlich, & Branco, S. (2020). Gene Copy Number Variation Does Not Reflect Structure or Environmental Selection in Two Recently Diverged California Populations of Suillus brevipes. G3 Genes|Genomes|Genetics, 10(12), 4591–4597. 10.1534/g3.120.401735

Beichman, A. C., Huerta-Sanchez, E., & Lohmueller, K. E. (2018). Using Genomic Data to Infer Historic Population Dynamics of Nonmodel Organisms. *Annual Review of Ecology*, Evolution, and Systematics, 49(1), 433–456. 10.1146/annurev-ecolsys-110617-062431

Behrens, C., Maciejewski, M. F., Arredondo, E., Dalziel, A. C., Weir, L. K., & Bell, A. M. (2024). Divergence in Reproductive Behaviors Is Associated with the Evolutionary Loss of Parental Care. The American Naturalist, 203(5), 590–603. 10.1086/729465

Behrens, C., Young, S., Arredondo, E., Dalziel, A. C., Weir, L. K., & Bell, A. M. (2025). The Evolutionary Loss of Paternal Care Is Associated With Shifts in Female Life-History Traits. Ecology and Evolution, 15(4), e70497. 10.1002/ece3.70497

Behrens, C., Maciejewski, M. F., Sumarli, A., Lucas, G., Lagunas-Robles, G., Samuk, K., & Bell, A. M. (2025). Evidence of parental care as a novel reproductive isolating mechanism. bioRxiv, 2025.04.18.649590. 10.1101/2025.04.18.649590

Bell, M. A., & Foster, S. A. (1994). Introduction to the evolutionary biology of the threespine stickleback. In M. A. Bell & S. a. Foster (Eds.), The Evolutionary Biology of the Threespine Stickleback (p. 0). Oxford University Press. 10.1093/oso/9780198577287.003.0001

Bell, S. E., Xie, Y. R., Maciejewski, M. F., Rubakhin, S. S., Romanova, E. V., Bell, A. M., & Sweedler, J. V. (2025). Single-Cell Peptide Profiling to Distinguish Stickleback Ecotypes with Divergent Breeding Behavior. Journal of Proteome Research, 24(4), 1596–1605. 10.1021/acs.jproteome.4c00832

Blackman, B. K. (2016). Speciation Genes. In Encyclopedia of Evolutionary Biology (pp. 166–175). Elsevier. 10.1016/B978-0-12-800049-6.00066-4

Blouw, D. M., & Hagen, D. W. (1990). Breeding ecology and evidence of reproductive isolation of a widespread stickleback fish (Gasterosteidae) in Nova Scotia, Canada. Biological Journal of the Linnean Society, 39(3), 195–217. 10.1111/j.1095-8312.1990.tb00512.x

Blouw, D. M. (1996). Evolution of offspring desertion in a stickleback fish. Écoscience, 3(1), 18–24. 10.1080/11956860.1996.11682310

Bolnick, D. I., Shim, K. C., & Brock, C. D. (2015). Female stickleback prefer shallow males: Sexual selection on nest microhabitat. Evolution, 69(6), 1643–1653. 10.1111/evo.12682

Chain, F. J. J., Feulner, P. G. D., Panchal, M., Eizaguirre, C., Samonte, I. E., Kalbe, M., Lenz, T. L., Stoll, M., Bornberg-Bauer, E., Milinski, M., & Reusch, T. B. H. (2014). Extensive Copy-Number Variation of Young Genes across Stickleback Populations. PLOS Genetics, 10(12), e1004830. 10.1371/journal.pgen.1004830

Cingolani, P., Platts, A., Wang, L. L., Coon, M., Nguyen, T., Wang, L., Land, S. J., Lu, X., & Ruden, D. M. (2012). A program for annotating and predicting the effects of single nucleotide polymorphisms, SnpEff: SNPs in the genome of Drosophila melanogaster strain w^1118^ ; iso-2; iso-3. Fly, 6(2), 80–92. 10.4161/fly.19695

Conrad, D. F., Pinto, D., Redon, R., Feuk, L., Gokcumen, O., Zhang, Y., Aerts, J., Andrews, T. D., Barnes, C., Campbell, P., Fitzgerald, T., Hu, M., Ihm, C. H., Kristiansson, K., MacArthur, D. G., MacDonald, J. R., Onyiah, I., Pang, A. W. C., Robson, S., … Hurles, M. E. (2010). Origins and functional impact of copy number variation in the human genome. Nature, 464(7289), 704–712. 10.1038/nature08516

Conte, G. L., & Schluter, D. (2013). Experimental Confirmation That Body Size Determines Mate Preference Via Phenotype Matching in a Stickleback Species Pair. Evolution, 67(5), 1477–1484. 10.1111/evo.12041

Corney, R. H., & Weir, L. K. (2023). Does paternal care influence mate preference? Male and female mating behavior in Threespine Stickleback ecotypes that differ markedly in parental care. Ecology and Evolution, 13(3), e9953. 10.1002/ece3.9953

Cotter, D. J., Brotman, S. M., & Wilson Sayres, M. A. (2016). Genetic Diversity on the Human X Chromosome Does Not Support a Strict Pseudoautosomal Boundary. Genetics, 203(1), 485–492. 10.1534/genetics.114.172692

Coughlan, J. M., & Matute, D. R. (2020). The importance of intrinsic postzygotic barriers throughout the speciation process. Philosophical Transactions of the Royal Society B: Biological Sciences, 375(1806), 20190533. 10.1098/rstb.2019.0533

Coyne, J. A., Coyne, H. A., & Orr, H. A. (2004). Speciation. Oxford University Press, Incorporated. https://books.google.com/books?id=Hq9RswEACAAJ

Cruickshank, T. E., & Hahn, M. W. (2014). Reanalysis suggests that genomic islands of speciation are due to reduced diversity, not reduced gene flow. Molecular Ecology, 23(13), 3133–3157. 10.1111/mec.12796

Cummings, M. E., & Endler, J. A. (2018). 25 Years of sensory drive: The evidence and its watery bias. Current Zoology, 64(4), 471–484. 10.1093/cz/zoy043

Czech, L., & Exposito-Alonso, M. (2022). grenepipe: A flexible, scalable and reproducible pipeline to automate variant calling from sequence reads. Bioinformatics, 38(20), 4809– 4811. 10.1093/bioinformatics/btac600

Danecek, P., Bonfield, J. K., Liddle, J., Marshall, J., Ohan, V., Pollard, M. O., Whitwham, A., Keane, T., McCarthy, S. A., Davies, R. M., & Li, H. (2021). Twelve years of SAMtools and BCFtools. GigaScience, 10(2), giab008. 10.1093/gigascience/giab008

Dean, L. L., Dunstan, H. R., Reddish, A., & MacColl, A. D. C. (2021). Courtship behavior, nesting microhabitat, and assortative mating in sympatric stickleback species pairs. Ecology and Evolution, 11(4), 1741–1755. 10.1002/ece3.7164

Delmore, K. E., Van Doren, B. M., Ullrich, K., Curk, T., van der Jeugd, H. P., & Liedvogel, M. (2023). Structural genomic variation and migratory behavior in a wild songbird. Evolution Letters, qrad040. 10.1093/evlett/qrad040

DeRaad, D. A. (2022). snpfiltr: An R package for interactive and reproducible SNP filtering. Molecular Ecology Resources, 22(6), 2443–2453. 10.1111/1755-0998.13618

Diniz, G. B., & Bittencourt, J. C. (2017). The Melanin-Concentrating Hormone as an Integrative Peptide Driving Motivated Behaviors. Frontiers in Systems Neuroscience, 11. 10.3389/fnsys.2017.00032

Dorant, Y., Cayuela, H., Wellband, K., Laporte, M., Rougemont, Q., Mérot, C., Normandeau, E., Rochette, R., & Bernatchez, L. (2020). Copy number variants outperform SNPs to reveal genotype–temperature association in a marine species. Molecular Ecology, 29(24), 4765– 4782. 10.1111/mec.15565

Dufresnes, C., & Crochet, P.-A. (2022). Sex chromosomes as supergenes of speciation: Why amphibians defy the rules? Philosophical Transactions of the Royal Society B: Biological Sciences, 377(1856), 20210202. 10.1098/rstb.2021.0202

Durinck, S., Spellman, P. T., Birney, E., & Huber, W. (2009). Mapping identifiers for the integration of genomic datasets with the R/Bioconductor package biomaRt. Nature Protocols, 4(8), 1184–1191. 10.1038/nprot.2009.97

Elphinstone, M. S., Hinten, G. N., Anderson, M. J., & Nock, C. J. (2003). An inexpensive and high-throughput procedure to extract and purify total genomic DNA for population studies. Molecular Ecology Notes, 3(2), 317–320. 10.1046/j.1471-8286.2003.00397.x

Emlen, S. T., & Oring, L. W. (1977). Ecology, Sexual Selection, and the Evolution of Mating Systems. Science, 197(4300), 215–223. 10.1126/science.327542

Endler, J. A. (1992). Signals, Signal Conditions, and the Direction of Evolution. The American Naturalist, 139, S125–S153.

Eskova, A., Frohnhöfer, H. G., Nüsslein-Volhard, C., & Irion, U. (2020). Galanin Signaling in the Brain Regulates Color Pattern Formation in Zebrafish. Current Biology, 30(2), 298–303.e3. 10.1016/j.cub.2019.11.033

Fang, B., Merilä, J., Ribeiro, F., Alexandre, C. M., & Momigliano, P. (2018). Worldwide phylogeny of three-spined sticklebacks. Molecular Phylogenetics and Evolution, 127, 613–625. 10.1016/j.ympev.2018.06.008

Feder, J. L., Nosil, P., Wacholder, A. C., Egan, S. P., Berlocher, S. H., & Flaxman, S. M. (2014). Genome-Wide Congealing and Rapid Transitions across the Speciation Continuum during Speciation with Gene Flow. Journal of Heredity, 105(S1), 810–820. 10.1093/jhered/esu038

Feller, A. F., Seehausen, O., Lucek, K. J. O., & Marques, D. A. (2024). Habitat Choice and Female Preference in a Polymorphic Stickleback Population. 10.7892/BORIS.79067

Gavrilets, S. (2000). Rapid evolution of reproductive barriers driven by sexual conflict. Nature, 403(6772), 886–889. 10.1038/35002564

Grant, V. J. (1993). Correlated Responses to Embryo Abandonment in a Newly Described Stickleback. Thesis. Antigonish, NS: St Francis Xavier University.

Gutenkunst, R. N., Hernandez, R. D., Williamson, S. H., & Bustamante, C. D. (2009). Inferring the Joint Demographic History of Multiple Populations from Multidimensional SNP Frequency Data. PLOS Genetics, 5(10), e1000695. 10.1371/journal.pgen.1000695

Haines, G. E. (2023). Intraspecific diversity of threespine stickleback (Gasterosteus aculeatus) populations in eastern Canada. Environmental Biology of Fishes, 106(5), 1177–1194. 10.1007/s10641-022-01362-1

Haley, A. L., Dalziel, A. C., & Weir, L. K. (2019). A comparison of nuptial coloration and breeding behaviour in white and common marine threespine stickleback (Gasterosteus aculeatus) ecotypes. Evolutionary Ecology Research, 20(2), 145–166.

Huang, K., Andrew, R. L., Owens, G. L., Ostevik, K. L., & Rieseberg, L. H. (2020). Multiple chromosomal inversions contribute to adaptive divergence of a dune sunflower ecotype. Molecular Ecology, 29(14), 2535–2549. 10.1111/mec.15428

Iskow, R. C., Gokcumen, O., & Lee, C. (2012). Exploring the role of copy number variants in human adaptation. Trends in Genetics : TIG, 28(6), 245–257. 10.1016/j.tig.2012.03.002

Irwin, D. E. (2018). Sex chromosomes and speciation in birds and other ZW systems. Molecular Ecology, 27(19), 3831–3851. 10.1111/mec.14537

Jamieson, I. G., Blouw, D. M., & Colgan, P. W. (1992). Parental care as a constraint on male mating success in fishes: A comparative study of threespine and white sticklebacks. Canadian Journal of Zoology, 70(5), 956–962. 10.1139/z92-136

Jamieson, I. G., Blouw, D. M., & Colgan, P. W. (1992). Field observations on the reproductive biology of a newly discovered stickleback (Gasterosteus). Canadian Journal of Zoology, 70(5), 1057–1063. 10.1139/z92-148

Jones, F. C., Grabherr, M. G., Chan, Y. F., Russell, P., Mauceli, E., Johnson, J., Swofford, R., Pirun, M., Zody, M. C., White, S., Birney, E., Searle, S., Schmutz, J., Grimwood, J., Dickson, M. C., Myers, R. M., Miller, C. T., Summers, B. R., Knecht, A. K., … Kingsley, D. M. (2012). The genomic basis of adaptive evolution in threespine sticklebacks. Nature, 484(7392), 55–61. 10.1038/nature10944

Kaplan, H. S., Horvath, P. M., Rahman, M. M., & Dulac, C. (2025). The neurobiology of parenting and infant-evoked aggression. Physiological Reviews, 105(1), 315–381. 10.1152/physrev.00036.2023

Kirkpatrick, M., & Barton, N. (2006). Chromosome inversions, local adaptation and speciation. Genetics, 173(1), 419–434. 10.1534/genetics.105.047985

Kokko, H., & Jennions, M. D. (2008). Parental investment, sexual selection and sex ratios. Journal of Evolutionary Biology, 21(4), 919–948. 10.1111/j.1420-9101.2008.01540.x

Korunes, K. L., & Samuk, K. (2021). pixy: Unbiased estimation of nucleotide diversity and divergence in the presence of missing data. Molecular Ecology Resources, 21(4), 1359– 1368. 10.1111/1755-0998.13326

Li, H., & Durbin, R. (2011). Inference of human population history from individual whole-genome sequences. Nature, 475(7357), 493–496. 10.1038/nature10231

Li, H., & Durbin, R. (2009). Fast and accurate short read alignment with Burrows-Wheeler transform. *Bioinformatics (Oxford*, England), 25(14), 1754–1760. 10.1093/bioinformatics/btp324

Liu, S., Hansen, M. M., & Jacobsen, M. W. (2016). Region-wide and ecotype-specific differences in demographic histories of threespine stickleback populations, estimated from whole genome sequences. Molecular Ecology, 25(20), 5187–5202. 10.1111/mec.13827

Maciejewski, M. F., Fischer, E. K., & Bell, A. M. (2025). An Evolutionary Loss of Parental Care in Stickleback Is Associated with Differences in the Activity, but Not the Number, of Neuropeptidergic Neurons in the Preoptic Area. Brain Behavior and Evolution. 10.1159/000545350

Marques, D. A., Lucek, K., Haesler, M. P., Feller, A. F., Meier, J. I., Wagner, C. E., Excoffier, L., & Seehausen, O. (2017). Genomic landscape of early ecological speciation initiated by selection on nuptial colour. Molecular Ecology, 26(1), 7–24. 10.1111/mec.13774

Mather, N., Traves, S. M., & Ho, S. Y. W. (2019). A practical introduction to sequentially Markovian coalescent methods for estimating demographic history from genomic data. Ecology and Evolution, 10(1), 579–589. 10.1002/ece3.5888

McKinnon, J. S., & Rundle, H. D. (2002). Speciation in nature: The threespine stickleback model systems. Trends in Ecology & Evolution, 17(10), 480–488. 10.1016/S0169-5347(02)02579-X

Meisel, R. P., & Connallon, T. (2013). The faster-X effect: Integrating theory and data. Trends in Genetics, 29(9), 537–544. 10.1016/j.tig.2013.05.009

Milinski, M., Griffiths, S., Wegner, K. M., Reusch, T. B., Haas-Assenbaum, A., & Boehm, T. (2005). Mate choice decisions of stickleback females predictably modified by MHC peptide ligands. Proceedings of the National Academy of Sciences of the United States of America, 102(12), 4414–4418. 10.1073/pnas.0408264102

Moehring, A. J., Llopart, A., Elwyn, S., Coyne, J. A., & Mackay, T. F. C. (2006). The Genetic Basis of Prezygotic Reproductive Isolation Between Drosophila santomea and D. yakuba Due to Mating Preference. Genetics, 173(1), 215–223. 10.1534/genetics.105.052993

Momigliano, P., Florin, A.-B., & Merilä, J. (2021). Biases in Demographic Modeling Affect Our Understanding of Recent Divergence. Molecular Biology and Evolution, 38(7), 2967– 2985. 10.1093/molbev/msab047

Nath, S., Shaw, D. E., & White, M. A. (2021). Improved contiguity of the threespine stickleback genome using long-read sequencing. G3 Genes|Genomes|Genetics, 11(2), jkab007. 10.1093/g3journal/jkab007

Nei, M., & Li, W. H. (1979). Mathematical model for studying genetic variation in terms of restriction endonucleases. Proceedings of the National Academy of Sciences of the United States of America, 76(10), 5269–5273. 10.1073/pnas.76.10.5269

Noor, M. a. F., & Bennett, S. M. (2009). Islands of speciation or mirages in the desert? Examining the role of restricted recombination in maintaining species. Heredity, 103(6), Article 6. 10.1038/hdy.2009.151

North, H. L., Caminade, P., Severac, D., Belkhir, K., & Smadja, C. M. (2020). The role of copy-number variation in the reinforcement of sexual isolation between the two European subspecies of the house mouse. Philosophical Transactions of the Royal Society B: Biological Sciences, 375(1806), 20190540. 10.1098/rstb.2019.0540

Ólafsdóttir, G. Á., Ritchie, M. G., & Snorrason, S. S. (2006). Positive assortative mating between recently described sympatric morphs of Icelandic sticklebacks. Biology Letters, 2(2), 250–252. 10.1098/rsbl.2006.0456

Paudel, Y., Madsen, O., Megens, H.-J., Frantz, L. A., Bosse, M., Bastiaansen, J. W., Crooijmans, R. P., & Groenen, M. A. (2013). Evolutionary dynamics of copy number variation in pig genomes in the context of adaptation and domestication. BMC Genomics, 14(1), 449. 10.1186/1471-2164-14-449

Pedersen, B. S., Layer, R., & Quinlan, A. R. (2020). smoove: structural-variant calling and genotyping with existing tools (Version 0.2.8) [Computer software]

Peichel, C. L., McCann, S. R., Ross, J. A., Naftaly, A. F. S., Urton, J. R., Cech, J. N., Grimwood, J., Schmutz, J., Myers, R. M., Kingsley, D. M., & White, M. A. (2020). Assembly of the threespine stickleback Y chromosome reveals convergent signatures of sex chromosome evolution. Genome Biology, 21(1), 177. 10.1186/s13059-020-02097-x

Perry, G. H., Dominy, N. J., Claw, K. G., Lee, A. S., Fiegler, H., Redon, R., Werner, J., Villanea, F. A., Mountain, J. L., Misra, R., Carter, N. P., Lee, C., & Stone, A. C. (2007). Diet and the evolution of human amylase gene copy number variation. Nature Genetics, 39(10), 1256–1260. 10.1038/ng2123

Picard Toolkit.” 2019. Broad Institute, GitHub Repository. https://broadinstitute.github.io/picard/; Broad Institute.

Poplin, R., Ruano-Rubio, V., DePristo, M. A., Fennell, T. J., Carneiro, M. O., Van der Auwera, G. A., Kling, D. E., Gauthier, L. D., Levy-Moonshine, A., Roazen, D., Shakir, K., Thibault, J., Chandran, S., Whelan, C., Lek, M., Gabriel, S., Daly, M. J., Neale, B., MacArthur, D. G., & Banks, E. (2018). Scaling accurate genetic variant discovery to tens of thousands of samples. bioRxiv, 201178. 10.1101/201178

Presgraves, D. C. (2010). The molecular evolutionary basis of species formation. Nature Reviews Genetics, 11(3), 175–180. 10.1038/nrg2718

Rafferty, N. E., & Boughman, J. W. (2006). Olfactory mate recognition in a sympatric species pair of three-spined sticklebacks. Behavioral Ecology, 17(6), 965–970. 10.1093/beheco/arl030

Raudvere, U., Kolberg, L., Kuzmin, I., Arak, T., Adler, P., Peterson, H., & Vilo, J. (2019). g:Profiler: A web server for functional enrichment analysis and conversions of gene lists (2019 update). Nucleic Acids Research, 47(W1), W191–W198. 10.1093/nar/gkz369

Ravinet, M., Faria, R., Butlin, R. K., Galindo, J., Bierne, N., Rafajlović, M., Noor, M. a. F., Mehlig, B., & Westram, A. M. (2017). Interpreting the genomic landscape of speciation: A road map for finding barriers to gene flow. Journal of Evolutionary Biology, 30(8), 1450–1477. 10.1111/jeb.13047

Redon, R., Ishikawa, S., Fitch, K. R., Feuk, L., Perry, G. H., Andrews, T. D., Fiegler, H., Shapero, M. H., Carson, A. R., Chen, W., Cho, E. K., Dallaire, S., Freeman, J. L., González, J. R., Gratacòs, M., Huang, J., Kalaitzopoulos, D., Komura, D., MacDonald, J. R., … Hurles, M. E. (2006). Global variation in copy number in the human genome. Nature, 444(7118), 444–454. 10.1038/nature05329

Reid, N. M., Proestou, D. A., Clark, B. W., Warren, W. C., Colbourne, J. K., Shaw, J. R., Karchner, S. I., Hahn, M. E., Nacci, D., Oleksiak, M. F., Crawford, D. L., & Whitehead, A. (2016). The genomic landscape of rapid repeated evolutionary adaptation to toxic pollution in wild fish. Science (New York, N.Y.), 354(6317), 1305–1308. 10.1126/science.aah4993

Reid, K., Bell, M. A., & Veeramah, K. R. (2021). Threespine Stickleback: A Model System For Evolutionary Genomics. Annual Review of Genomics and Human Genetics, 22(Volume 22, 2021), 357–383. 10.1146/annurev-genom-111720-081402

Rieseberg, L. H. (2001). Chromosomal rearrangements and speciation. Trends in Ecology & Evolution, 16(7), 351–358. 10.1016/s0169-5347(01)02187-5

Roesti, M., Gavrilets, S., Hendry, A. P., Salzburger, W., & Berner, D. (2014). The genomic signature of parallel adaptation from shared genetic variation. Molecular Ecology, 23(16), 3944–3956. 10.1111/mec.12720

Rougeux, C., Bernatchez, L., & Gagnaire, P.-A. (2017). Modeling the Multiple Facets of Speciation-with-Gene-Flow toward Inferring the Divergence History of Lake Whitefish Species Pairs (Coregonus clupeaformis). Genome Biology and Evolution, 9(8), 2057– 2074. 10.1093/gbe/evx150

Sambrook, J., & Russell, D. W. (2006). Purification of nucleic acids by extraction with phenol:chloroform. CSH protocols, 2006(1), pdb.prot4455. 10.1101/pdb.prot4455

Samuk, K. M. (2016). *The evolutionary genomics of adaptation and speciation in the threespine stickleback* (T). University of British Columbia. Retrieved from https://open.library.ubc.ca/collections/ubctheses/24/items/1.0307145

Sardell, J. M., Josephson, M. P., Dalziel, A. C., Peichel, C. L., & Kirkpatrick, M. (2021). Heterogeneous Histories of Recombination Suppression on Stickleback Sex Chromosomes. Molecular Biology and Evolution, 38(10), 4403–4418. 10.1093/molbev/msab179

Schield, D. R., Scordato, E. S. C., Smith, C. C. R., Carter, J. K., Cherkaoui, S. I., Gombobaatar, S., Hajib, S., Hanane, S., Hund, A. K., Koyama, K., Liang, W., Liu, Y., Magri, N., Rubtsov, A., Sheta, B., Turbek, S. P., Wilkins, M. R., Yu, L., & Safran, R. J. (2021). Sex-linked genetic diversity and differentiation in a globally distributed avian species complex. Molecular Ecology, 30(10), 2313–2332. 10.1111/mec.15885

Schluter, D., & McPhail, J. D. (1992). Ecological Character Displacement and Speciation in Sticklebacks. The American Naturalist. 10.1086/285404

Schluter, D., & Rieseberg, L. H. (2022). Three problems in the genetics of speciation by selection. Proceedings of the National Academy of Sciences, 119(30), e2122153119. 10.1073/pnas.2122153119

Sendrowski, J., & Bataillon, T. (2024). fastDFE: Fast and Flexible Inference of the Distribution of Fitness Effects. Molecular Biology and Evolution, 41(5), msae070. 10.1093/molbev/msae070

Shaw, K. L., Parsons, Y. M., & Lesnick, S. C. (n.d.). QTL analysis of a rapidly evolving speciation phenotype in the Hawaiian cricket Laupala. Retrieved April 25, 2025, from https://onlinelibrary.wiley.com/doi/10.1111/j.1365-294X.2007.03321.x

Stankowski, S., Cutter, A. D., Satokangas, I., Lerch, B. A., Rolland, J., Smadja, C. M., Segami Marzal, J. C., Cooney, C. R., Feulner, P. G. D., Bicalho Domingos, F. M. C., North, H. L., Yamaguchi, R., Butlin, R. K., Wolf, J. B. W., Coughlan, J., Heidbreder, P., Hernández-Gutiérrez, R., Barnard-Kubow, K. B., Peede, D., … Kulmuni, J. (2024). Toward the integration of speciation research. Evolutionary Journal of the Linnean Society, kzae001. 10.1093/evolinnean/kzae001

Suvakov, M., Panda, A., Diesh, C., Holmes, I., & Abyzov, A. (2021). CNVpytor: A tool for copy number variation detection and analysis from read depth and allele imbalance in whole-genome sequencing. GigaScience, 10(11), giab074. 10.1093/gigascience/giab074

Sylvestre, F., Mérot, C., Normandeau, E., & Bernatchez, L. (2023). Searching for intralocus sexual conflicts in the three-spined stickleback (Gasterosteus aculeatus) genome. Evolution, 77(7), 1667–1681. 10.1093/evolut/qpad075

Therkildsen, N. O., & Palumbi, S. R. (2017). Practical low-coverage genomewide sequencing of hundreds of individually barcoded samples for population and evolutionary genomics in nonmodel species. Molecular ecology resources, 17(2), 194–208. 10.1111/1755-0998.12593

Thompson, K. A., Brandvain, Y., Coughlan, J. M., Delmore, K. E., Justen, H., Linnen, C. R., Ortiz-Barrientos, D., Rushworth, C. A., Schneemann, H., Schumer, M., & Stelkens, R. (2024). The Ecology of Hybrid Incompatibilities. Cold Spring Harbor Perspectives in Biology, 16(9), a041440. 10.1101/cshperspect.a041440

Turbek, S. P., Browne, M., Di Giacomo, A. S., Kopuchian, C., Hochachka, W. M., Estalles, C., Lijtmaer, D. A., Tubaro, P. L., Silveira, L. F., Lovette, I. J., Safran, R. J., Taylor, S. A., & Campagna, L. (2021). Rapid speciation via the evolution of pre-mating isolation in the Iberá Seedeater. Science, 371(6536), eabc0256. 10.1126/science.abc0256

Van der Auwera, G. A., Carneiro, M. O., Hartl, C., Poplin, R., Del Angel, G., Levy-Moonshine, A., Jordan, T., Shakir, K., Roazen, D., Thibault, J., Banks, E., Garimella, K. V., Altshuler, D., Gabriel, S., & DePristo, M. A. (2013). From FastQ data to high confidence variant calls: The Genome Analysis Toolkit best practices pipeline. Current Protocols in Bioinformatics, 43(1110), 11.10.1-11.10.33. 10.1002/0471250953.bi1110s43

Via, S. (2009). Natural selection in action during speciation. Proceedings of the National Academy of Sciences, 106(supplement_1), 9939–9946. 10.1073/pnas.0901397106

Webster, T. H., & Wilson Sayres, M. A. (2016). Genomic signatures of sex-biased demography: Progress and prospects. Current Opinion in Genetics & Development, 41, 62–71. 10.1016/j.gde.2016.08.002

Weir, B. S., & Cockerham, C. C. (1984). Estimating F-Statistics for the Analysis of Population Structure. Evolution, 38(6), 1358–1370.JSTOR. 10.2307/2408641

Wilson Sayres, M. A. (2018). Genetic Diversity on the Sex Chromosomes. Genome Biology and Evolution, 10(4), 1064–1078. 10.1093/gbe/evy039

Wu, Z., Autry, A. E., Bergan, J. F., Watabe-Uchida, M., & Dulac, C. G. (2014). Galanin neurons in the medial preoptic area govern parental behavior. Nature, 509(7500), 325–330. 10.1038/nature13307

Yi, X., Liang, Y., Huerta-Sanchez, E., Jin, X., Cuo, Z. X. P., Pool, J. E., Xu, X., Jiang, H., Vinckenbosch, N., Korneliussen, T. S., Zheng, H., Liu, T., He, W., Li, K., Luo, R., Nie, X., Wu, H., Zhao, M., Cao, H., … Wang, J. (2010). Sequencing of 50 human exomes reveals adaptation to high altitude. Science (New York, N.Y.), 329(5987), 75–78. 10.1126/science.1190371

Yilmaz, O., Sullivan, C. V., Bobe, J., & Norberg, B. (2024). The role of multiple vitellogenins in early development of fishes. General and Comparative Endocrinology, 351, 114479. 10.1016/j.ygcen.2024.114479

Zhang, Y., Hu, Y., Wang, X., Jiang, Q., Zhao, H., Wang, J., Ju, Z., Yang, L., Gao, Y., Wei, X., Bai, J., Zhou, Y., & Huang, J. (2020). Population Structure, and Selection Signatures Underlying High-Altitude Adaptation Inferred From Genome-Wide Copy Number Variations in Chinese Indigenous Cattle. Frontiers in genetics, 10, 1404. 10.3389/fgene.2019.01404

Zhao, H., Sun, Z., Wang, J., Huang, H., Kocher, J.-P., & Wang, L. (2014). CrossMap: A versatile tool for coordinate conversion between genome assemblies. Bioinformatics, 30(7), 1006–1007. 10.1093/bioinformatics/btt730

Zheng, X., Levine, D., Shen, J., Gogarten, S. M., Laurie, C., & Weir, B. S. (2012). A high-performance computing toolset for relatedness and principal component analysis of SNP data. Bioinformatics, 28(24), 3326–3328. 10.1093/bioinformatics/bts606

