## Supplementary Materials (Low Coverage Library Preparation, Figures S1, S2, Tables S3-S5 for "Strong but diffuse genetic divergence underlies differentiation in an incipient species of marine stickleback"

**Low Coverage Library Preparation**

Quantified samples were laid out in a 384-well PCR plate and molecular grade water was used as a negative control for the library preparation. For each 2.5 µL reaction, 1 µL of normalized DNA was added along with 1.25 µL TD buffer and 0.25 µL tagmentation enzyme. Tagmentation reactions were run at 55°C for 5 mins followed by a 10°C hold.

Following tagmentation, all individual libraries were indexed in 7.5 µL PCRs composed of 3.76 µL KAPA HiFi 2X Master Mix (Roche Diagnostics, Rotkreuz, Switzerland) and 0.,625 µL of both N5 and N7 Nextera combinatorial indices at 5 µM concentration (Illumina, San Diego USA). Reactions were run with the following protocol: 72°C for 3 min, 98°C for 2 min 45 s, (98°C for 15 s, 62°C for 30 s, 72°C for 1 min 30 s) x 8, 72°C for 1 min 30 s, 10°C hold.

Following indexing, a reconditioning PCR using the Illumina adapter sequences as primers was done to further enrich indexed library fragments. For each indexing reaction, 9.5 µL of reconditioning PCR mix composed of 8.5 µL KAPA HiFI 2X Master Mix, and 0.5 µL of each primer at 10 µM was added directly to the previous PCR product. The reconditioning PCR was run with the following parameters: 95°C for 5 min, (98°C for 10 s, 62°C for 20 s, 72°C for 30 s) x 4, 72°C for 2 minutes, 4°C hold. For pooling, 4 µL of each PCR was pipetted into a 1.5 mL Low Bind microcentrifuge tube (Sarstedt, Nümbrecht, Germany).

Pooled libraries were cleaned using a one-sided bead clean protocol. A 1:1 ratio of Illumina SPB beads and pooled library were combined and mixed, incubated at room temperature for 15 min, and placed on a magnet for an additional 5 minutes. Following this, the supernatant was decanted and the beads were washed twice with freshly prepared 80% ethanol for 30 seconds before decanting. Beads were then air-dried on the magnet for 5 minutes before being removed. To elute the cleaned library, 25µL of Tris-Tween (10mM tris, 0.05% Tween-20) was added and mixed with the beads, incubated at room temperature for 2 minutes, placed on the magnet for 5 min, and ~25 µL of eluted library was transferred to a new 1.5 ml Low Bind microcentrifuge tube for storage. A second bead clean was performed to further remove primer dimer.

Successful library preparation was confirmed by assessing a 10K dilution of the cleaned library via qPCR using a KAPA 2X Library Quantification Kit with primers added (Roche Diagnostics). Reactions were composed of 4.5 µL of KAPA 2X Library Quant Master Mix with primers added, 1 µL molecular-grade water, and 2 µL of standard, diluted library, or molecular-grade water for the blank. The qPCR was performed following manufacturer protocols and concentrations were calculated assuming library fragments were generally 2x larger than the Illumina standards.

**Figure S1.** PCA of SNPs from 20 Common and 20 White sticklebacks from CB and CL respectively. **A.** PCA of 806,908 autosomal SNPs, **B.** PCA of 49,480 SNPs from the X chromosome in females. **C.** PCA of 11,411 SNPs from the Y chromosome in males.

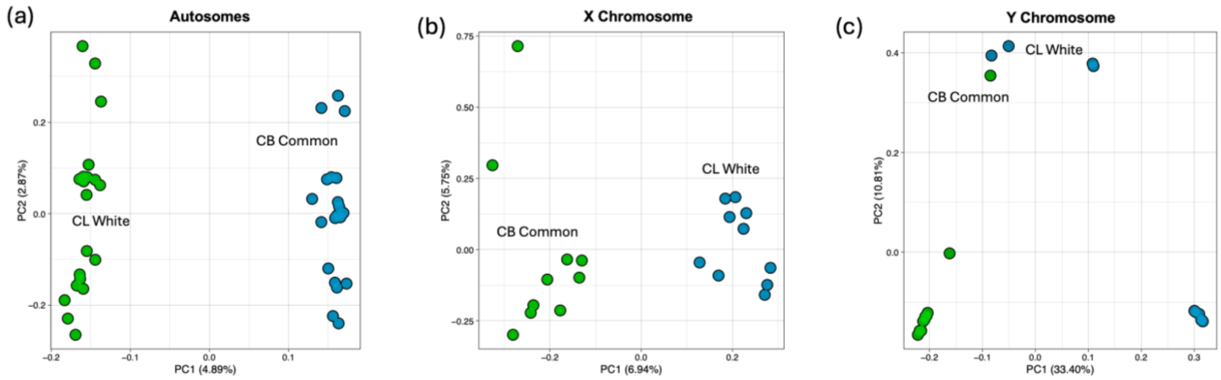


**Figure S2.** (a) Plot of $F_{ST}$ on y-axis and $V_{ST}$ on x-axis of cross-validated deletions and duplications. Regression line in red. (b) Size distribution of cross-validated duplications with dotted lines representing mean length. (c) Distance of cross-validated duplications from genes with dotted lines representing mean length. Values of 0 indicate CNVs overlap with genes.

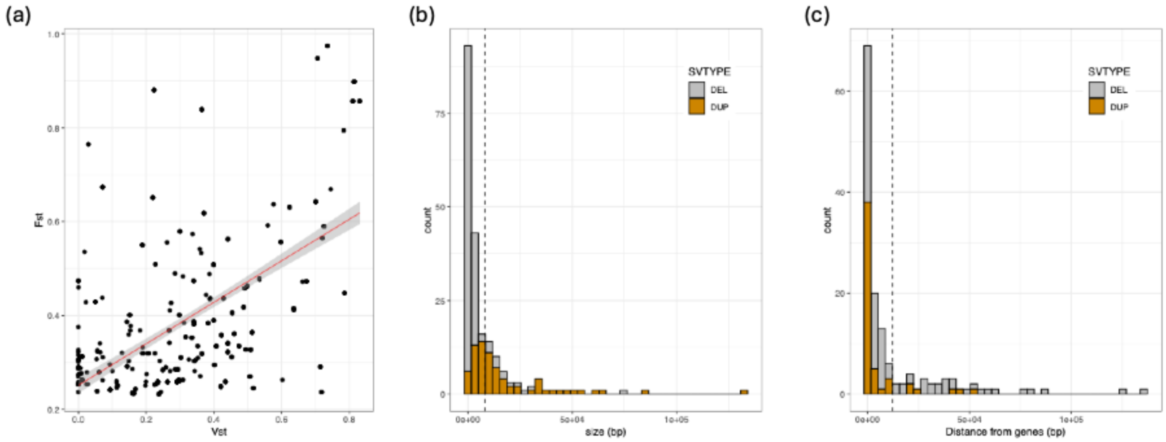


**Table S1.** Information about each individual analyzed in this study including sex, ecomorph, location, and accession number. Chromosomes analyzed (i.e., autosome, X chromosome, Y chromosome) are demarcated by "+".

**Table S2.** Demographic model parameters and scaled parameters estimated by dadi-cli for all eleven models tested. The top three models with confidence intervals and residuals are included on the second page. All symbols follow Table 2.

**Table S3.** Population Branch Statistic of each population. Mean and upper and lower limits of the 95% confidence interval are shown for each summary statistic. 95% confidence intervals were calculated from 1,000 bootstrap replicates.

| **Population** | **Autosome PBS** | **X Chromosome PBS** | **Y Chromosome PBS** |
|---|---|---|---|
| White | 0.0263 ± 0.0001 | 0.0295 ± 0.0006 | 0.1772 ± 0.0157 |
| Nova Scotia Common | 0.0102 ± 0.0001 | 0.0206 ± 0.0005 | 0.1602 ± 0.0131 |
| Quebec Outgroup | 0.0176 ± 0.0001 | 0.0296 ± 0.0008 | 0.0317 ± 0.0053 |

**Table S4.** Pairwise Weir and Cockerham's $F_{ST}$ of each population pair used to calculate branch lengths for the PBS analysis. Mean and upper and lower limits of the 95% confidence interval are shown for each summary statistic. 95% confidence intervals were calculated from 1,000 bootstrap replicates.

| **Population Pair** | **Autosome $F_{ST}$** | **X Chromosome $F_{ST}$** | **Y Chromosome $F_{ST}$** |
|---|---|---|---|
| White - Common | 0.0244 ± 0.0001 | 0.0331 ± 0.0006 | 0.2666 ± 0.0081 |
| White - Quebec | 0.0332 ± 0.0001 | 0.0426 ± 0.0008 | 0.2541 ± 0.0071 |
| Quebec - Common | 0.0162 ± 0.0001 | 0.0326 ± 0.0007 | 0.1795 ± 0.0062 |

**Table S5.** $F_{ST}$ and $V_{ST}$ of deletions and duplications on the autosomes, X Chromosome, and Y Chromosome of white and commons. Mean, SD, and upper and lower limits of the 95% confidence interval are shown for each summary statistic. 95% confidence intervals were calculated from 1,000 bootstrap replicates

| **Statistic** | **Autosome** | **X Chromosome** | **Y Chromosome** |
|---|---|---|---|
| CNV $F_{ST}$ | 0.016 ± 0.0004 | 0.0208 ± 0.0024 | 0.2064 ± 0.0405 |

| | | | |
|---|---|---|---|
| CNV $V_{ST}$ | 0.0324 ± 0.0006 | 0.0427 ± 0.0036 | 0.0801 ± 0.0148 |

**Table S6.** White PBS outliers on the autosomes and X chromosome with GO terms and enrichment analysis results.

**Table S7.** CNV outliers on the autosomes and X chromosome. Includes GO terms and enrichment analysis results.
